# Genomic heterogeneity and ploidy identify patients with intrinsic resistance to PD-1 blockade in metastatic melanoma

**DOI:** 10.1101/2022.12.11.519808

**Authors:** Giuseppe Tarantino, Cora A. Ricker, Annette Wang, Will Ge, Tyler J. Aprati, Amy Y. Huang, Shariq Madha, Jiajia Chen, Yingxiao Shi, Marc Glettig, Dennie T. Frederick, Samuel Freeman, Marta M. Holovatska, Michael P. Manos, Lisa Zimmer, Alexander Rösch, Anne Zaremba, Brendan Reardon, Jihye Park, Haitham A. Elmarakeby, Bastian Schilling, Anita Giobbie-Hurder, Natalie I. Vokes, Elizabeth I. Buchbinder, Keith T. Flaherty, Rizwan Haq, Catherine J. Wu, Genevieve M. Boland, F. Stephen Hodi, Eliezer M. Van Allen, Dirk Schadendorf, David Liu

## Abstract

While the introduction of immune checkpoint blockade (ICB) has dramatically improved clinical outcomes for patients with advanced melanoma, a significant proportion of patients develop resistance to therapy, and mechanisms of resistance are poorly elucidated in most cases. Further, while combination ICB has higher response rates and improved progression free survival compared to single agent therapy in the front line setting, there is significantly increased toxicity with combination ICB, and biomarkers to identify patients who would disproportionately benefit from combination therapy vs aPD-1 ICB are poorly characterized. To understand resistance mechanisms to single vs combination ICB therapy, we analyze whole-exome-sequencing (WES) of pre-treatment tumor and matched normals of 4 cohorts (n=140) of previously ICB-naïve aPD-1 ICB treated patients. We find that high intratumoral genomic heterogeneity and low ploidy identify patients with intrinsic resistance to aPD-1 ICB. Comparing to a melanoma cohort from a pre-targeted therapy and ICB time period (“untreated” cohort), we find that genomic heterogeneity specifically predicts response and survival in the ICB treated cohorts, but not in the untreated cohort, while ploidy is also prognostic of overall survival in the “untreated” (by targeted therapy or ICB) group. To establish clinically actionable predictions, we optimize a simple decision tree using genomic ploidy and heterogeneity to identify with high confidence (90% PPV) a subset of patients with intrinsic resistance to and significantly worse survival on aPD1 ICB treatment. We then validate this model in independent cohorts, and further show that a significant proportion of patients predicted to have intrinsic resistance to single agent aPD-1 ICB respond to combination ICB, which suggests that nominated patients may benefit disproportionately from combination ICB. We further show that the features and predictions of the model are independent of known clinical features and previously nominated molecular biomarkers. These findings highlight the clinical and biological importance of genomic heterogeneity and ploidy, and sets a concrete framework towards clinical actionability, broadly advancing precision medicine in oncology.

## Introduction

The introduction of immune checkpoint blockade (ICB) has dramatically improved the treatment landscape for patients with advanced melanoma, but only a subset of patients have durable response to therapy[^1–3^]. Single agent aPD1 ICB nivolumab and pembrolizumab as well as combination aPD1/CTLA4 ICB (ipilimumab + nivolumab) are standard first-line treatment options for patients with advanced metastatic melanoma, with combination aPD1/CTLA4 ICB demonstrating improved response rates, PFS, and a strong trend towards improved OS compared to single agent aPD1 ICB[^4^]. However, combination therapy has a much higher rate of severe immune-related adverse events (>50% vs ∼15% for single agent aPD1 ICB) [^3,5^], while the absolute difference in proportion of patients with durable response to combination vs single agent ICB is < 10%. Thus, biomarkers to identify patients who would disproportionately benefit from combination versus single agent ICB in the front line setting would reduce toxicity while optimizing disease-specific outcomes. Currently no molecular biomarkers have been well-validated to guide these treatment decisions. This highlights the need to improve our understanding of the molecular determinants of response and resistance to (1) guide more personalized and rational utilization of ICB treatment options and (2) identify novel targets and combinations to overcome resistance. Thus far, several markers have been suggested to be associated with response to aPD-1 ICB. Tumor mutational burden (TMB) was the first to be associated with response in melanoma patients [^6,7^]. Subsequently, several additional features have been proposed based on neoantigen load, immunohistochemical quantification of PD-L1 and CD8, genetic alteration in the antigen presentation genes and gene expression-based IFN-γ signature [^7–13^]. Many of these biomarkers were nominated in non-melanoma or pan-cancer settings, with inconsistent validation in metastatic melanoma and without differentiation of important clinical context (e.g. different ICB regimens or prior therapy). In recent work predicting response to aPD1 ICB, we found that prior therapy was a significant stratifier and different features were associated with therapy response in patients with and without prior treatment with aCTLA4 ICB. We developed parsimonious predictive models integrating clinical and molecular features, but were limited in our ability to validate these models due to lack of available independent cohorts with the requisite data [^14^]. In this study, we focus on aCTLA4 ICB naïve metastatic melanoma patients (which represents the current front-line therapy setting for metastatic melanoma) treated with aPD-1 ICB, finding that genomic heterogeneity and ploidy predict intrinsic resistance to aPD-1 ICB, and refine our understanding of their predictive (under therapy) vs prognostic (independent of therapy) role in response and survival. We develop a simple modified decision tree based on these features to identify with high precision patients with intrinsic resistance to aPD-1 ICB who may disproportionately benefit from combination ICB, and validate these findings in independent cohorts of PD-1 and contrast with combination PD-1/CTLA4 ICB treated melanoma patients. We further find that genomic heterogeneity and ploidy, and the predictions of our model for patients with intrinsic resistance to PD-1 ICB, did not reflect known clinical features or previously nominated molecular features associated with poor-risk disease or poor response to ICB.

## Results

### Low ploidy and high heterogeneity discriminate patients with intrinsic resistance to aPD-1 ICB in multiple independent cohorts

We harmonized several metastatic melanoma patients cohorts described in previous studies and clinical trials (supplementary table 1) [^15,16^], focusing on the subset of patients that were previously ICB-naïve, treated with aPD1 ICB, and had available WES data of pre-treatment tumor samples (Methods) to identify patients with intrinsic resistance to therapy (progressive disease (PD) at first restaging, hereafter also referred to as “progressors”). In a previous integrated genomic, transcriptomic, and clinical analysis of metastatic melanoma patients treated with aPD1 ICB ^14^, we found that a logistic regression model with features of genomic heterogeneity, ploidy, and tumor purity predicted intrinsic resistance to aPD1 ICB in previously ICB-naïve patients. In an independent cohort of patients from two clinical trials (BMS CheckMate-038 and −064 (respectively, 27 and 13 patients included after QC)), the nominated model had a modest AUC of 0.64 (Supplementary figure 1A; AUC in the original cohort 0.76), but examining the individual features of the model, tumor purity was not associated with response in the independent cohorts evaluated (Supplementary figure 1B), but the association between higher genomic heterogeneity and lower ploidy with intrinsic resistance to therapy was robust (ploidy MW p=0.002, heterogeneity MW p=0.038; ploidy MW p=0.027, heterogeneity MW p=0.018 in the original and independent cohorts, respectively) (Figure 1A and B). Thus, we developed new models using only genomic heterogeneity and ploidy in a new combined discovery cohort (Figure 1C) of n=124 patients. Both logistic regression and decision tree models using heterogeneity and ploidy had moderate AUCs in the combined cohort (AUC of 0.73 and 0.75, respectively; 10 fold cross-validation AUC 0.72; Supplementary figure 2). A prediction of PD was associated with an ORR of 5.1 [95% CI 2.4-10.9] and 8.6 [95% CI 3.7-20.1] for the logistic regression and decision tree models, respectively (Fig 1D &E). These models also stratified overall survival (OS) and progression free survival (PFS) with the patients predicted as PD possessing worse OS (Decision tree HR=3.1 [95% CI 1.8-5.3], p < 0.0001; logistic regression HR=1.9 [95% CI 1.1-3.3], p=0.019) and PFS (Supplementary figure 3; Decision tree HR=2.5 [95% CI 1.6-3.9], p < 0.0001; logistic regression HR=2.0 [95% CI 1.3-3.1], p=0.0018). The decision tree model was characterized by higher precision/positive predictive value (76% vs 66%) and specificity (84% vs 71%) compared to the logistic regression model (Supplemental Fig 2B and C), and provided a straightforward approach to predicting patients with intrinsic resistance (Fig 1F). Overall, we found that high genomic heterogeneity and low ploidy was robustly associated with intrinsic resistance to aPD1 therapy in multiple cohorts, and simple predictive models using these two features identified patients with intrinsic resistance with reasonable performance.

**Figure1.**
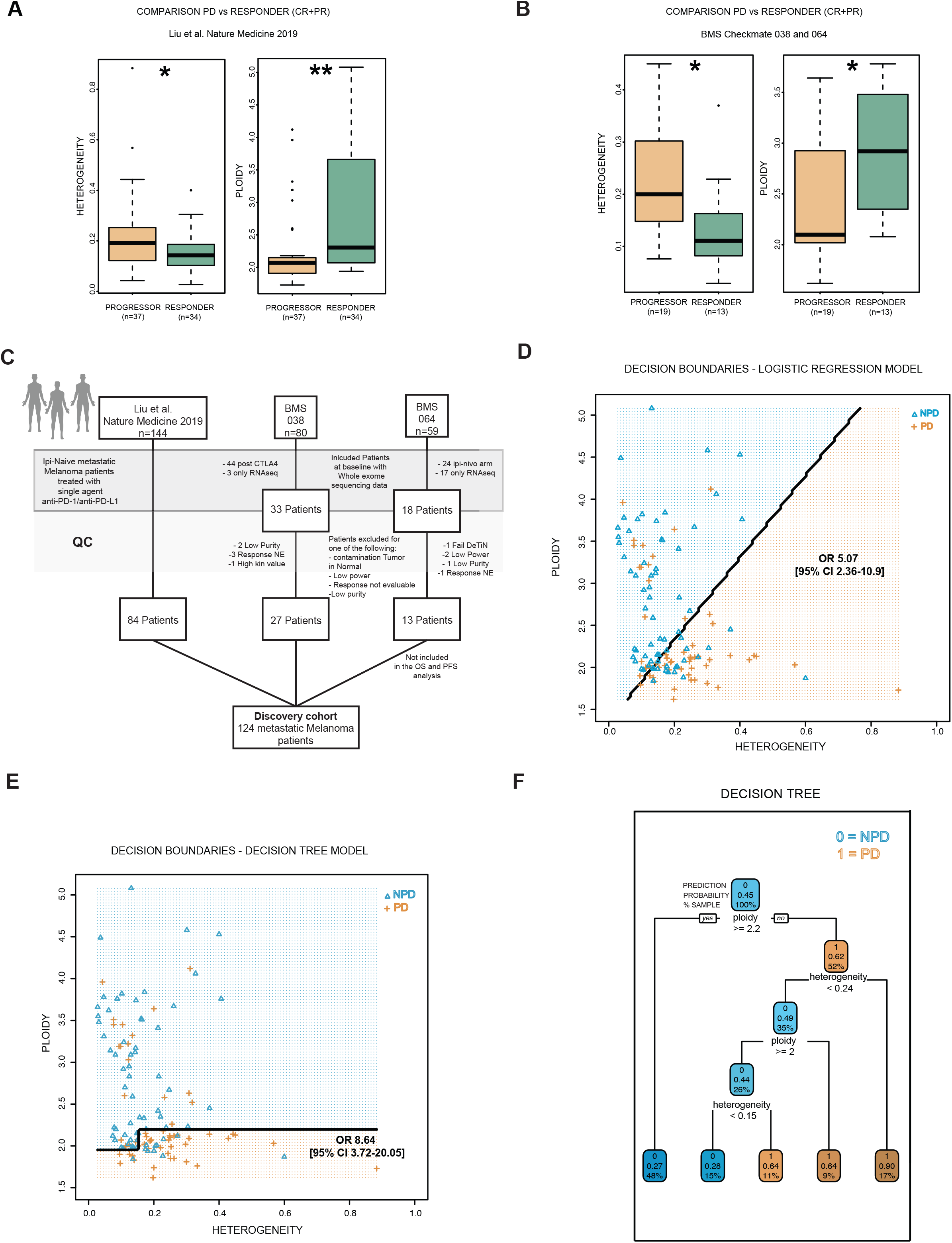
High genomic heterogeneity and low ploidy predict intrinsic resistance in previously ICB-naïve PD-1 treated patients. **A**. Genomic heterogeneity and ploidy in progressors (PD as best response, orange) vs responders (CR/PR as best response, green) in the CTLA-4 ICB naïve PD-1 ICB treated subset of a large discovery cohort of metastatic melanoma patients. (MWW p = 0.038, p=0.0021 for heterogeneity, ploidy, respectively) **B**. Genomic heterogeneity and ploidy comparison in progressors vs responders of a validation CTLA-4 ICB naïve PD-1 ICB treated cohort drawn from two clinical trials (MWW p = 0.018, p=0.027 for heterogeneity, ploidy, respectively. **C**. A new combined discovery cohort was constructed combining the patients from three different cohorts: the Liu et al. Nature Medicine 2019 paper, the clinical trials Checkmate 038 and 064. Checkmate-064 was a trial of sequential ipilimumab-nivolumab vs nivolumab-ipilimumab, and only the patients in the arm A (treated first with nivolumab) were selected; for these patients the response was evaluated after 12 weeks, and their data was not included in the survival analysis presented in this work. **D**. Decision boundaries of the logistic regression model (LR) with genomic heterogeneity and ploidy as features to predict patients with intrinsic resistance (PD) using the new combined discovery cohort. The orange area represents the area predicted by the model as PD while the blue area represents the patients predicted as non-progressive disease (nPD). The observed therapy response of each patient is represented by the orange plus symbol (PD) or the blue triangle (nPD). **E**. Decision boundaries (similar to D) for a decision tree model (DT). **F**. Structure of the decision tree with logic and split cutoff used. In each node: the top number represents the overall prediction for the node with 1 = PD and 0 = nPD; the second number represents the probability of the patients in that group to be PD; the third number denotes the proportion of samples in that node.

### Timing of WGD event distinguishes responders versus nonresponders with WGD

We analyzed tumors misclassified by our model, e.g. tumors with low heterogeneity and high ploidy but observed to have intrinsic resistance, and the converse. Higher ploidy in tumors is associated with response to aPD-1 ICB in our data (Supp Fig 4) and is driven by whole genome doubling (WGD) events (Figure 2A and B). Mutations within WGD tumors have different multiplicity (i.e. one or two copies per cancer cell) representing mutations that occurred after (1 copy) or before (2 copies) the WGD event (Figure 2C). The ratio of 2:1 multiplicity of mutations is thus associated with time from the WGD event [^17,18^]. Interestingly, 3 WGD tumors misclassified by our predictive model as non-progressors based on low heterogeneity and high ploidy had high 2:1 SNV multiplicity ratio, suggesting that they may represent recent WGD events (Fig 2D). Including SNV multiplicity as a feature of the model led to a small AUC improvement (Supp Fig 5A & B, with examples of PD samples with EGD and low heterogeneity in panel C; and Responders samples with WGD and low heterogeneity in panel D). Conversely, misclassified patients predicted to have intrinsic resistance (PD) but observed to have non-progressive disease (nPD) did not have distinguishable genomic or clinical features (Supplementary table 3). Most of these patients had stable disease as best response (7 SD, 3 PR, 1 CR out of 11 misclassified patients), and most misclassifications occurred at relatively lower heterogeneity (Supplementary Fig 6), suggesting poorer outcomes even if not progressive disease at the earliest time point.

**Figure2.**
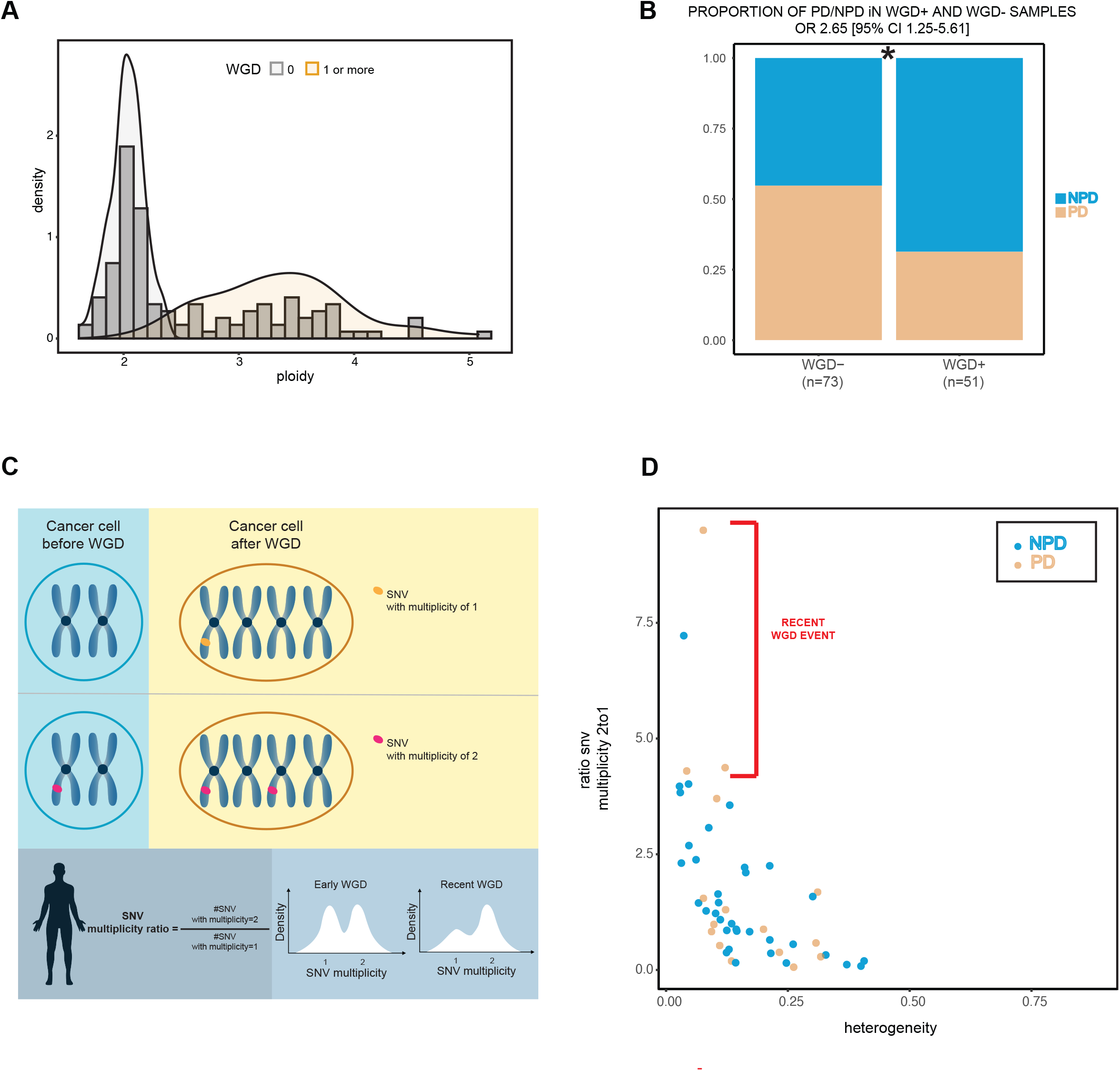
Whole-genome doubling and its timing is associated with response to ICB. **A**. Ploidy distribution of whole-genome doubled (WGD) and non-whole-genome-doubled tumors. Higher ploidy is driven by WGD events. **B**. Proportion of patients with WGD event in the PD patients vs NPD patients (Fisher’s exact p = 0.011). **C**. Graphical representation on how to compute the SNV multiplicity ratio and estimate the time of WGD event. **D**. Ratio of multiplicity 2:1 SNV mutations and heterogeneity scatterplot for WGD tumors. Orange dots represent patients with PD as best response (PD), and blue NPD. A high 2:1 SNV multiplicity ratio indicates few SNV mutations after genome doubling, consistent with a recent WGD event.

### Optimizing a predictive model to identify patients with intrinsic resistance with high specificity

To establish a clinically actionable predictive model, we developed a model to identify patients with intrinsic resistance to aPD-1 ICB prioritizing high specificity (i.e. high precision/positive predictive value (PPV)) over sensitivity (identifying all patients with intrinsic resistance correctly), reasoning that it may be clinically useful to identify patients with high probability of intrinsic resistance to single agent aPD-1 ICB who may disproportionately benefit from combination immunotherapies. Accordingly, we developed a modified version of the decision tree model (MDT) (online methods) using heterogeneity and ploidy (Figure 3A and Supplementary figure 6B). Using this model, 21 patients (17% of the cohort) were predicted to be PD, with a PPV of 90% (19/21 correctly predicted) and specificity of 97% (66/68 patients correctly identified as nPD). The models stratified overall survival (OS) and progression free survival (PFS) with the patients predicted as PD possessing worse OS (MDT HR=3.0 [95% CI 1.6-5.5], p = 0.00023) and PFS (Supplementary figure 6C; Decision tree HR=3.0 [95% CI 1.7-5.2], p < 0.0001).

**Figure3.**
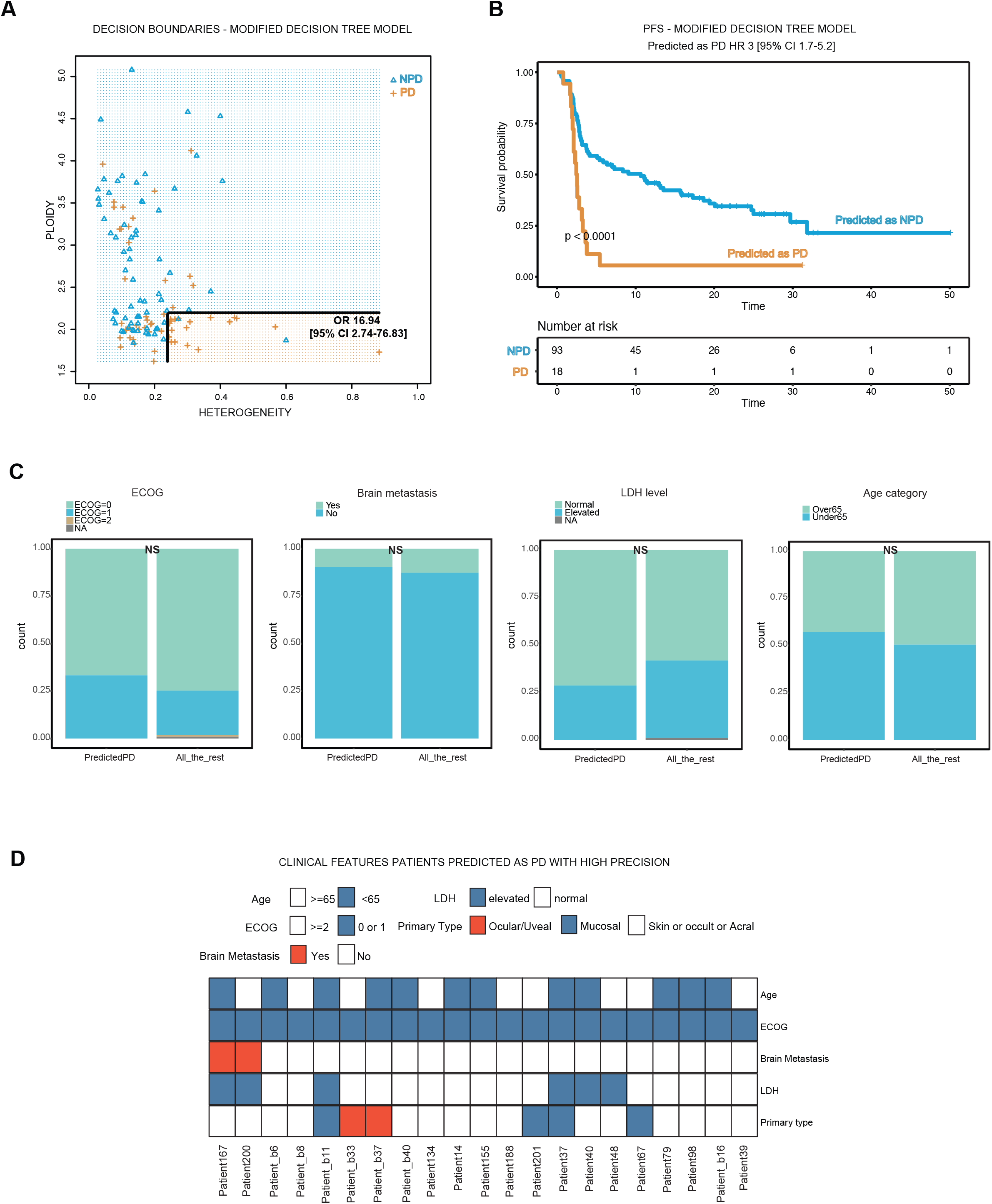
Constructing a modified decision tree model optimizing precision and specificity for predicting intrinsic resistance to PD-1 ICB. **A**. Decision boundaries of the modified decision tree model (MDT). **B**. Progression free survival curve stratified by patients predicted by the MDT model as PD (orange) and NPD (blue) (log rank p < 0.0001); in the survival analysis have been excluded the samples from Checkmate 064 that received a sequential treatment (n=13). **C**. Clinical characteristics between predicted PD patients (n=21) and the rest of the cohort, comparing ECOG, presence of Brain metastasis, LDH level at baseline and Age category; pvalues from fisher exact test. **D**. Clinical features of the 21 patients predicted as PD; only 4 patients (highlighted in red) have clinical features (brain metastasis, ocular/uveal primary type) that strongly indicate combination ICB.

### Patients predicted as intrinsically resistance to aPD-1 ICB have similar clinical features to the overall cohort

Certain clinical characteristics (e.g. high tumor burden, site-specific metastases (e.g. brain, liver) denoting worse disease, uveal melanoma) are associated with worse outcomes and often prompt treatment with combination ICB [^19,20,21^]. To understand whether patients predicted to be intrinsically resistant to aPD-1 ICB by our model also have worse clinical characteristics, we evaluated M stage, LDH baseline level, presence of brain met, primary melanoma subtype, presence of liver metastasis, ECOG, presence of lung metastasis and age. Overall, we found no statistically significant difference in clinical characteristics in patients predicted to be PD vs others in the cohort (Figure 3C). Further, out of the 21 patients predicted as PD only 4 possessed any known clinical features (2 with brain metastases, 2 with uveal melanoma) that would strongly favor the choice of combination ICB (Figure 3D, Supplementary table 2). Interestingly, the non-cutaneous melanoma subtypes had higher heterogeneity compared to cutaneous melanomas (acral (n=9, p=0.021), Ocular/Uveal patients (n=2,p=0.025), Mucosal group (n=9, p<0.001))(Supplementary figure 8). However, heterogeneity and ploidy were associated with response even when limiting the analysis to cutaneous melanoma (heterogeneity pval=0.013, ploidy pval<0.001; Supplementary figure 8D). We did not identify significant differences in heterogeneity and ploidy in terms of M stage, LDH level and between patients with and without brain metastasis (Supplementary figure 9 and 10). Overall, our analysis suggested that our model predicted patients with intrinsic resistance who otherwise did not have other clinical features that would have suggested more aggressive disease or resistance to aPD-1 ICB.

### Our model can better define PD and nPD patients compared to established genomic signatures and biomarkers

Tumor mutational burden and an interferon gamma signature have been associated with response to aPD-1 ICB in large pan-cancer cohorts [^10^], though their performance in melanoma-specific cohorts is uneven (AUC 0.60 and 0.64 for TMB and IFN-γ signature, respectively). In this cohort of metastatic melanoma patients, we tested the stratification of responders (CR+PR) vs PD in terms of TMB and IFN-γ. In the studies of the individual cohorts in our combined discovery cohort, the association of TMB with response to therapy was mixed[^14^]. Indeed, in our combined discovery cohort, one of the highest TMB tumors (>50 mut/MB) was a non-responder, but had high heterogeneity and low ploidy (Supplementary figure 11) and was correctly classified by our model. While TMB is independent of heterogeneity and ploidy (Supplementary Figure 7), adding TMB to the feature space does not significantly improve performance (TMB in the logistic regression model p=0.08) and is not supported by an AIC/BIC metric (used to trade off improvement in model performance with increased complexity of the model) (e.g. BIC increase of 1.74). For a subgroup of patients for whom the RNAseq data were available (n=108, Supplementary figure 12), IFN-γ was not correlated with ploidy and heterogeneity but does not improve model performance (Supplementary Figure 13). Finally, we evaluated a recently developed clinical nomogram [^22^] predicting response to ICB based on clinical features; unfortunately, not all the samples in our cohort had available clinical features used by the model (i.e. neutrophil to lymphocyte ratio, liver metastasis presence, ECOG, and lung metastasis presence). However, in 5 patients we had sufficient available clinical data to determine that they would be estimated by the nomogram to be at least intermediate or good response risks, but due to their low ploidy and high heterogeneity our model correctly predicted them to have intrinsic resistance (Supplementary figure 14), suggesting additional predictive information being provided by this genomic data.

### Genomic heterogeneity specifically predicts therapy response, while ploidy is prognostic

To understand the prognostic (i.e. indicating poor biology independent of therapy) vs predictive (outcome in the setting of therapy) roles of genomic heterogeneity and ploidy, we analyzed data from a TCGA melanoma cohort which was collected in a time frame where modern targeted and ICB therapies were not widely available (the “untreated” cohort). Ploidy was significantly higher in metastatic (n=392) vs primary (n=61) lesions (Fig 4A) consistent with past studies[^14,23,24^], showing WGD involvement in tumor evolution and metastasis [^25^]. In contrast, genomic heterogeneity was significantly higher in primary samples (Fig 4A), consistent with a founder bottlenecking effect in metastatic lesions. In univariate Cox survival analyses of the metastatic subset, ploidy but not heterogeneity was associated with overall survival (Figure 4C-D; heterogeneity HR = 1.5 [95% CI 0.55-4.0], p = 0.44, ploidy HR=0.76 [95% CI 0.6-0.96], p=0.02). In contrast, in our aPD-1 ICB treated cohort, high heterogeneity in metastatic samples was strongly associated with worse PFS and OS (PFS Cox HR = 8.0 [95% CI 1.5-42], p = 0.013; OS Cox HR = 19.0 [95% CI 4.1-84]), while ploidy had similar (but borderline statistically significant) associations with improved PFS (Cox HR = 0.74 [95% CI 0.54-1], p = 0.065) and OS (Cox HR = 0.76 [95% CI 0.51-1.1], p = 0.181) in this smaller cohort. Notably, the effect-size estimates of ploidy on survival was similar between the untreated and PD-1 ICB treated cohorts, but were not statistically significant in the aPD-1 ICB treated cohort potentially due to smaller sample size. In the multivariate analysis, ploidy (but not heterogeneity) again predicted overall survival in the untreated cohort (Figure 4E; HR = 0.64 [95% CI 0.5-0.84], p = 0.001), while in the aPD-1 ICB treated cohort heterogeneity strongly stratified PFS and OS while ploidy was no longer a strong predictor (Figure 4F and G; heterogeneity HR = 13.87 [95% CI 2.7-71.3], p = 0.002; ploidy HR = 0.87 [95% CI 0.56-1.4], p=0.55). Taken together, our analysis demonstrates a strong predictive role of genomic heterogeneity on patient outcomes under aPD-1 ICB therapy but not in untreated patients, while ploidy is also prognostic in the non-ICB treated setting.

**Figure4.**
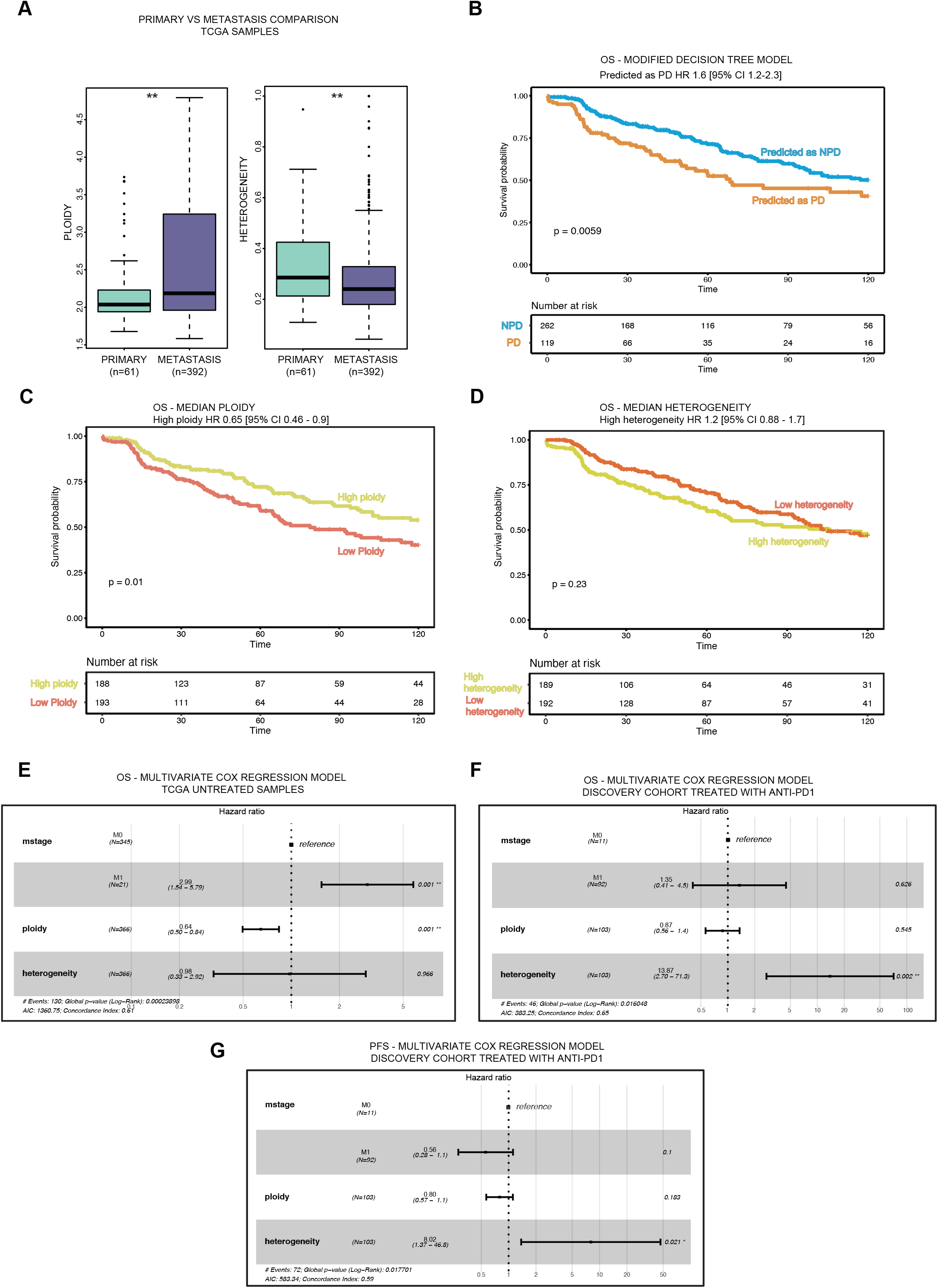
Association of heterogeneity and ploidy with survival in ICB-treated and -untreated cohorts. **A**. Difference in genomic heterogeneity and ploidy between primary and metastatic ICB-untreated samples in the TCGA melanoma cohort. (MWW p = 0.0031, p=0.0062 for heterogeneity and ploidy, respectively). **B**. OS survival of the TCGA samples stratified by predicted PD status using the modified DT model. (log rank p = 0.0059)**C**. OS survival of the TCGA samples stratified by median ploidy (log rank p=0.01). **D**. OS survival of the TCGA samples stratified by median heterogeneity (log rank p=0.23). **E**. Multivariate cox regression model evaluating the effect of ploidy and heterogeneity for the OS in the TCGA cohort. **F**. Multivariate cox regression model evaluating the effect of ploidy and heterogeneity for the OS in the anti-PD1 discovery cohort. **G**. Multivariate cox regression model evaluating the effect of ploidy and heterogeneity for the PFS in the anti-PD1 discovery cohort.

### Model validation in independent cohorts

Finally, to validate this model in an external cohort, we collected and tested our model against a small independent cohort of 16 additional patients who were ipilimumab-naïve treated with aPD1/aPD-L1 [^26^] with 4 patients with CR/PR, 1 patient with SD, 1 patient with mixed response (MR), and 10 patients with PD as BOR. Even in this small cohort of patients high heterogeneity and low ploidy identifies intrinsically resistant patients (Figure 5A-C). Further, our modified model continued to have high precision, with all patients predicted by our optimized model to be intrinsically resistant correctly predicted (n=5, PPV = 100% and specificity = 100%). We further applied our model to an independent cohort of combination aPD1/aCTLA4 ICB treated patients (n=13). Since RECIST annotation was not available for this cohort, we defined intrinsic resistance (PD) as patients who progressed with a PFS < 6 months vs the patients with PFS higher than 6 months (non-PD). Interestingly, heterogeneity still continued to have a trend towards being higher in PD patients vs non-PD (p = 0.052, Figure 5D); but for ploidy there was no significant difference. Strikingly, 3/7 (43%) of patients predicted to be PD to single agent PD-1 ICB in our model were non-PD when treated with combination aPD-1/aCTLA-4 ICB, suggesting that some of the patients identified by this model may differentially benefit from combination ICB compared to single agent ICB (Figure 5E).

**Figure5.**
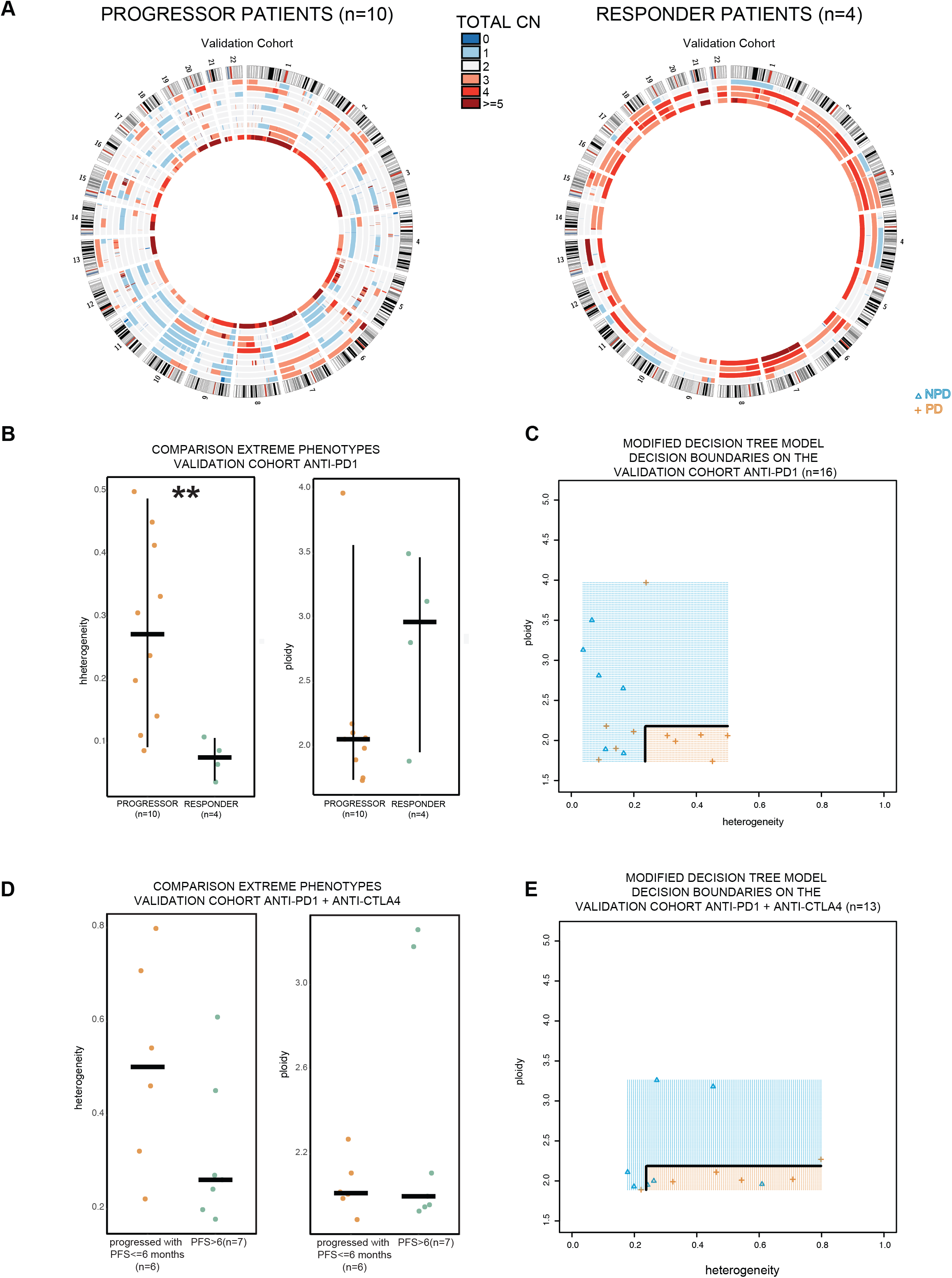
Association of heterogeneity, ploidy, and predicted PD-1 ICB intrinsic resistance with ICB response in independent validation cohorts. **A**. Circos plots of copy number alterations in progressors (PD as best response, left) and responders (CR/PR as best response, right) in a PD-1 ICB treated validation cohort. **B**. Heterogeneity and ploidy compared in progressors vs responders in the validation PD-1 ICB cohort (MWW p =0.008, p=0.23 for heterogeneity and ploidy, respectively). **C**. Decision boundaries for the modified decision tree model using the samples from the validation anti-PD1 ICB cohort. **D**. Heterogeneity and ploidy compared in responders (PFS > 6 months) vs progressors (PFS <= 6 months) in a combination PD-1/CTLA-4 ICB cohort (MWW p = 0.1, p=1 for heterogeneity and ploidy, respectively). **E**. Decision boundaries for the modified decision tree model using the samples from the combination PD-1/CTLA-4 ICB cohort showing response in 3/7 patients predicted to be intrinsically resistant to PD-1 ICB.

## Discussion

In this study, we identified low ploidy and high genomic heterogeneity as two robust independent biomarkers of intrinsic resistance to aPD1 ICB in metastatic melanoma patients without prior ICB in multiple independent cohorts. We then developed a simple predictive model using genomic heterogeneity and ploidy to identify with high precision a subset of patients with intrinsic resistance to single agent aPD1 ICB. Our results demonstrated that these patients do not possess other adverse clinical characteristics that would have indicated poor risk disease. We further identified genomic heterogeneity as uniquely predictive in the setting of ICB response, while ploidy is prognostic of overall survival in an untreated cohort of metastatic melanoma patients. Further, timing of whole-genome-doubling event, which is the primary driver of ploidy differences, may impact predictions of response and survival to aPD-1 ICB.

Intratumoral genomic heterogeneity has been associated with highly mutagenic disease with an higher likelihood of preexisting or rapidly evolving resistant clones, and associated with worse clinical outcomes in a range of clinical contexts[^14,27–30^]. *In vivo* studies of intratumoral heterogeneity using mixes of UVB-irradiated subclones have demonstrated increased tumor growth in heterogeneous compared to homogenous tumors in immune-competent vs immunodeficient mice [^31^]. However, it has variable association with response to ICB in different histologies and clinical contexts, and its utility as a biomarker has not previously been well-demonstrated. The role of ploidy is much more complex. Large-scale differences in ploidy are driven by WGD events, which are common in cancer (30% of solid tumors in one estimate[^23^]). The hypothesized benefit of WGD to tumors is the ability to tolerate copy number alterations (aneuploidy) across the genome to find more favorable genomic states [^25^], and mitigate the accumulation of deleterious somatic alterations [^32^]. Indeed, in this study we observe WGD and increased aneuploidy at later tumor stages (Fig 4A) and longitudinally in other studies in individual patients [^33^]. Further, the pro- or anti-tumorigenic effects of WGD may be context-dependent [^34–36^], where WGD is associated with tumorigenesis by increasing aneuploidy [^37–39^], but may also activate cellular stress mechanisms including the p53 and Hippo pathways, as well as immune surveillance [^37,40,41^]. Prior work has suggested that tumors with WGD may have unique vulnerabilities [^42^], but the mechanisms underlying the observation that WGD tumors are associated with ICB response (e.g. neoantigen presentation, increase in immunogenicity) are unclear. Interestingly, our observation that timing of WGD, as measured by SNV multiplicity ratio, may be associated with outlier resistance to ICB suggests that the increased vulnerability to ICB occurs later in WGD tumors. Clinically, our results suggest that SNV multiplicity should be evaluated together with WGD and aneuploidy in future studies of biomarkers of therapy response.

Interestingly, our data demonstrates that intratumoral genomic heterogeneity is specifically predictive of response and survival in a ICB therapy setting, while ploidy is also prognostic of worse outcomes (i.e. poor biology) even in a non-targeted or immuno-therapy treated setting. This suggests that clinical staging stratifying survival may be improved by incorporating molecular markers.

Importantly, we developed a novel approach to developing biomarkers by optimizing specificity and precision in predicting patients with intrinsic resistance to aPD-1 ICB. By design, we are tolerating reduced sensitivity (i.e. identification of all intrinsically resistant patients) for increased positive predictive value of the predicted resistant patients. Clinically, this translates into higher confidence of predictions in a smaller subset of patients, which may improve clinical applicability and adoption. Our model predicted intrinsic resistance in ∼20% of the entire cohort, with 90% PPV, and validated in a small independent cohort (5/5 patients correctly predicted to have intrinsic resistance). We further asked if our model simply replicates known genomic or clinical features of poor prognosis disease and response to ICB and found that our predictions were independent of known and nominated features and clinical nomograms, suggesting the independent utility of these predictions. Further, application of our model in a small cohort of combination aPD-1/aCTLA-4 ICB treated patients shows response in a significant subset (3/7, 43%) of patients predicted to have intrinsic resistance to aPD-1 ICB, suggesting that these are indeed patients who would disproportionately benefit from combination therapy in the front line setting.

There remain several limitations to this study. First, our independent validation cohorts were relatively small (in part due to careful curation of ICB-naïve tumor samples). Validation in larger cohorts is necessary, though our findings have been robust in every cohort so far. Second, our assays are based on biopsies of single lesions at a single point in time, which multiple studies[^43,44^] have demonstrated may not be accurate representations of tumor genomic heterogeneity even within the same lesion, and differences in biopsy sites may bias our estimates. However, within our limited data, we do not observe consistent differences between genomic heterogeneity and ploidy between biopsy sites (Supplementary Fig 15), and our results suggest high precision even without explicitly accounting for these differences. Third, the specific biological mechanisms underpinning the association of intratumoral heterogeneity and ploidy with ICB response (and prognosis) are unclear and represent important future research directions. Fourth, limiting clinical actionability, intratumoral heterogeneity and ploidy are estimated here using WES of matched tumor and normal tissue and associated analytics which are not standardized or routinely available clinically. Standardized assessments of these genomic metrics [^45^] remain to be developed using clinically validated assays in prospective settings. However, we have found our results to be robust using a different automated tool[^46^] to infer tumor heterogeneity and ploidy (Supplementary Fig 16), suggesting the feasibility of this approach.

Taken together, we have demonstrated genomic heterogeneity and ploidy as robust predictors of intrinsic resistance to aPD1 ICB in metastatic melanoma, clarified the predictive vs prognostic role of these features in metastatic melanoma, developed a novel approach to constructing clinically relevant predictive models, constructed and validated a new predictive model that identifies patients with intrinsic resistance to aPD1 ICB with high confidence, laying the foundation for prospective studies to translate these findings to clinical practice. Broadly, this represents a significant advance in the development and application of molecular biomarkers in precision oncology.

## Online Methods

### Patient cohorts

Metastatic melanoma patients treated with immune checkpoint blockade were identified from published work (Liu et al. Nature Medicine 2019 & Freeman et al. Cell Reports Medicine 2022[^14,26^]) and completed clinical trials (BMS Checkmate 038 and checkmate 064). We included only samples without prior exposure to ipilimumab, with WES data of the paired tumor and normal tissue obtained before PD1 blockade. Clinicopathological and demographic data were obtained from Liu et al Nature Medicine 2019, from BMS for the two clinical trials and for the validation cohort from Freeman et al. [^26^]. Data are shown in Figure 1 and in Supplementary table 1. The best objective response (BOR) to aPD1 ICB was only available for a subgroup of the patients included in Freeman et al. and wasn’t available for the combination immunotherapy-treated (“combo”) cohort. The analysis was performed by defining responders as patients achieving CR or PR as BOR; patients showing PD as BOR were instead defined as progressors. To understand if intrinsic resistant patients to aPD1 could benefit from the combo ICB, we also included a previously unpublished internal cohort of combo treated samples. For the combo cohort, for which the BOR was not available, we defined patients that progressed in the first six months of treatment as progressors and compared them versus patients with PFS > 6 months. The definition of OS and PFS was from initiation of ICB and sample collection as described in their respective studies.

This retrospective study and associated informed consent procedures were approved by the central Ethics Committee (EC) of the University Hospital Essen (12-5152-BO and 11-4715) and of Dana Farber Cancer Institute (IRB 05-042). Approval by the local EC was obtained by investigators if required by local regulations.

### Quality control and variant calling

Samples from the BMS and Freeman et al. cohorts were re-analyzed with the Broad institute CGA pipeline [^47–57^] using the TERRA platform, adopting the same quality controls filters used for the Liu et al. Nature Medicine 2019. In particular quality control cutoffs were as follows: mean target coverage > 50X (tumor) and >30X (normal), cross contamination of samples estimation (ContEst)<5%, tumor purity >= 10%, DeTiN ≤ 20% TiN. A power filter combining coverage and tumor purity was applied as described (e.g. minimum 80% power to detect clonal mutations) in Liu et al. Nature Medicine 2019. Three samples were excluded for low purity and two samples for low power.

MuTect2 [^47^] was used to identify somatic single-nucleotide variants in targeted exons, with computational filtering of artifacts introduced by DNA oxidation during sequencing or FFPE-based DNA extraction using a filter-based method [^50^]. Subsequently Strelka [^49^] was used to identify small insertions or deletions. Lastly, Oncotator [^57^] was used to annotate the Identified alterations.

### Ploidy, Purity and heterogeneity estimation

Absolute was used for the estimation of ploidy, purity and for the cancer cell fraction (CCF) estimation of individual mutations [^54^]. For each sample, the optimal solution (purity, ploidy) was manually selected among the local solutions. Heterogeneity was computed as the proportion of the subclonal mutations, with a mutation defined as subclonal if the cancer cell fraction (CCF) was lower than 0.8.

For the TCGA samples, ploidy and heterogeneity were taken from Conway et al. Nat Genet 2020 (which used FACETS^46^ to estimate purity, ploidy, and individual mutation CCF).

### SNV multiplicity and the time of WGD

The snv multiplicity for each SNV, representing the number of copies of the SNV per cancer cell, was estimated using tumor purity and the estimated copy number state at the SNV site (q_hat) from ABSOLUTE^54^ to estimate the expected variant allele fraction associated with a multiplicity of 1:

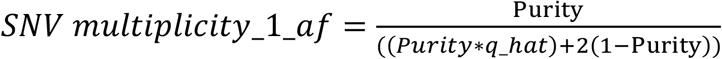

Then SNV multiplicity was estimated using the observed tumor variant allele fraction (Tumor_f) and the expected variant allele fraction for a multiplicity of 1.

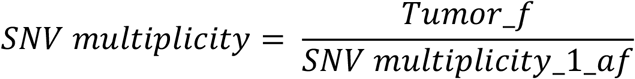

We then assigned each SNV to either multiplicity 1 or 2 based on a cutoff according to the distribution of the SNVs, selecting the lowest point in the histogram between the two modes of multiplicity at 1 and 2 in each individual sample (Supplementary table 1). For each sample, the ratio of SNV multiplicity 2:1 alterations was used as a metric of time since the WGD event; patients with a recent WGD are characterized by high SNV multiplicity 2:1 ratio [^17,18^]. In the modified logistic regression model the SNV multiplicity, since it is a feature of only the WGD samples, was included as:

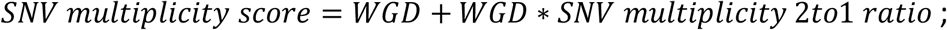

where WGD is 0 or 1, 1 for the samples with one or more WGD events.

#### Predictive model generation

In order to develop an interpretable predictive model, we focused on two model types; a logistic regression and a decision tree model. Both the models were based on just two features: heterogeneity and ploidy. The model was trained to predict PD as the best RECIST response versus non-PD (nPD) rather than responder versus progressor to better reflect the real-world setting where all outcomes (PD, SD, MR, PR and CR) are possible. We also evaluated the prediction accuracy of a logistic regression model including the snv multiplicity as additional feature. For the standard decision tree model, we used default complexity, method=“class”, and in order to avoid overfitting, we used the value of 10 as minimum number of samples included in the leaf. The modified decision tree model to optimize precision was obtained by increasing relative weight of nPD samples vs PD samples in a 4-fold cross validation procedure repeated 10 times (using R package *caret v 6*.*0*.*93*). The choice of relative weight (nPD = 2) for the final model was selected for a tradeoff between increased precision or PPV (elbow method) and decreased sensitivity (supplementary figure 17). The models were implemented in *R* version 4.2.0 using the packages *stats (v 4*.*2*.*0)* and *rpart (v 4*.*1*.*19)*; for the confusion matrix and the metrics of the models was used the R package *caret*. To estimate the cross-validation AUC of the logistic regression model, we used k-fold cross-validation using k=10 (splitting the dataset into k-subsets, training on k − 1 subsets and calculating AUC on the holdout subset) and calculated the mean cross-validation AUC and standard deviations. Cross-validation scores were calculated using the *cross_val_score* function from the Python (v3) *sklearn* (v 1.0.2) package.

### TCGA analysis

The TCGA data was obtained from Conway et al. Nat Genet 2020 [^58^]. Heterogeneity was calculated using the same cutoff used for the ABSOLUTE analysis (cancer cell fraction > 0.8 defined clonal mutations). Samples were initially divided into primary (n=61) and metastatic (n=392) samples to compare heterogeneity and ploidy; subsequent analyses focused on the metastatic lesions with OS data (n=381).

### Transcriptomic analysis

The methods used for sample collection, sequencing and quality control have been described in previous work [^14^]. For a subset of the samples included in the discovery cohort bulk RNAseq was available (n=108). Only transcriptomes from tumors whose WES also passed quality control were included.

To evaluate the role of the interferon γ, we used ssGSEA and signature genesets:

- IFN-γ and IFN-γ related from Rodig et al. Sci. Transl. Med. 2018^15^
- The HALLMARK_INTERFERON_GAMMA_RESPONSE^59^

we compared progressors and responders in the three cohorts used. The analysis was implemented in R using the package GSVA ^60^ (v 1.44.0) and msigdbr ^61^(v7.5.1).

### Statistics and Reproducibility

Statistical analyses were performed using the *stats R* package for R version 4.2.0. Reported p-values represent nominal p-values. Two primary response comparisons were made: (1) responders (defined as having CR or PR as the best RECIST response) versus progressors (defined as having PD as the best RECIST response) and (2) progressors (PD as the best RECIST response) versus non-progressors (non-PD as best RECIST response). For the comparison between continuous clinical and molecular features the Mann-Whitney test was used. For association of binary variables Fisher’s exact test was used. All statistical tests performed were two-sided.

### Survival Analysis

The survival outcome of patients receiving aPD-1 was evaluated with Kaplan-Meier survival analysis. The significance of the difference in survival outcome between the patients predicted as PD (stratified by the P(PD)>50% for the logistic regression) was assessed using a two-sided log-rank test from the *survival R* package. We performed this test for both overall survival (OS) and progression free survival (PFS). Checkmate 064 was a trial of sequential therapy with nivolumab and ipilimumab, therefore patients from this trial (n=13 with first-line aPD1 ICB) were only used to identify intrinsic resistance (PD at the first restaging scan) and were excluded for the survival analysis. For the survival analysis, the cohort evaluated is n=111 metastatic melanoma patients. The impact of clinical and molecular features on overall survival and progression free survival was also tested using univariate and multivariate Cox proportional hazards model using *R* version 4.2.0 and the packages *survival (v 3*.*3*.*1)* and *survminer (v 0*.*4*.*9)*.

## Code availability

Code to regenerate figures from the data provided with this study is available at github at https://github.com/davidliu-lab. Additional requests for code will be promptly reviewed by the senior authors to verify whether the request is subject to any intellectual property or confidentiality obligations and shared to the extent permissible by these obligations.

## Data availability

All analyzed data are in supplementary tables or data available on github at https://github.com/davidliu-lab. Data to reproduce the work of Liu et al. 2019, BMS Checkmate038 and 064 findings, and Freeman et al. cohort have been already published and are included in the supplementary table with the same labels [^14–16,26^]. Raw sequencing data of new samples included in this analysis are available in dbgap (accession number phs000452.v3.p1).

## Acknowledgments

This work was supported in part by the US National Institutes of Health (NIH) and the Doris Duke Charitable Foundation. This work was funded by Bristol Myers Squibb through its International Immuno-Oncology Network. G.T. was supported by an American-Italian Cancer Foundation Post-Doctoral Research Fellowship, year 1. The results presented here are in part based upon data generated by TCGA Research Network.

