## Supplementary figures for "Genomic heterogeneity and ploidy identify patients with intrinsic resistance to PD-1 blockade in metastatic melanoma"

**Supplementary Figure 1. Evaluation of previously nominated predictive model utilizing genomic heterogeneity, ploidy, and purity in validation cohort.**

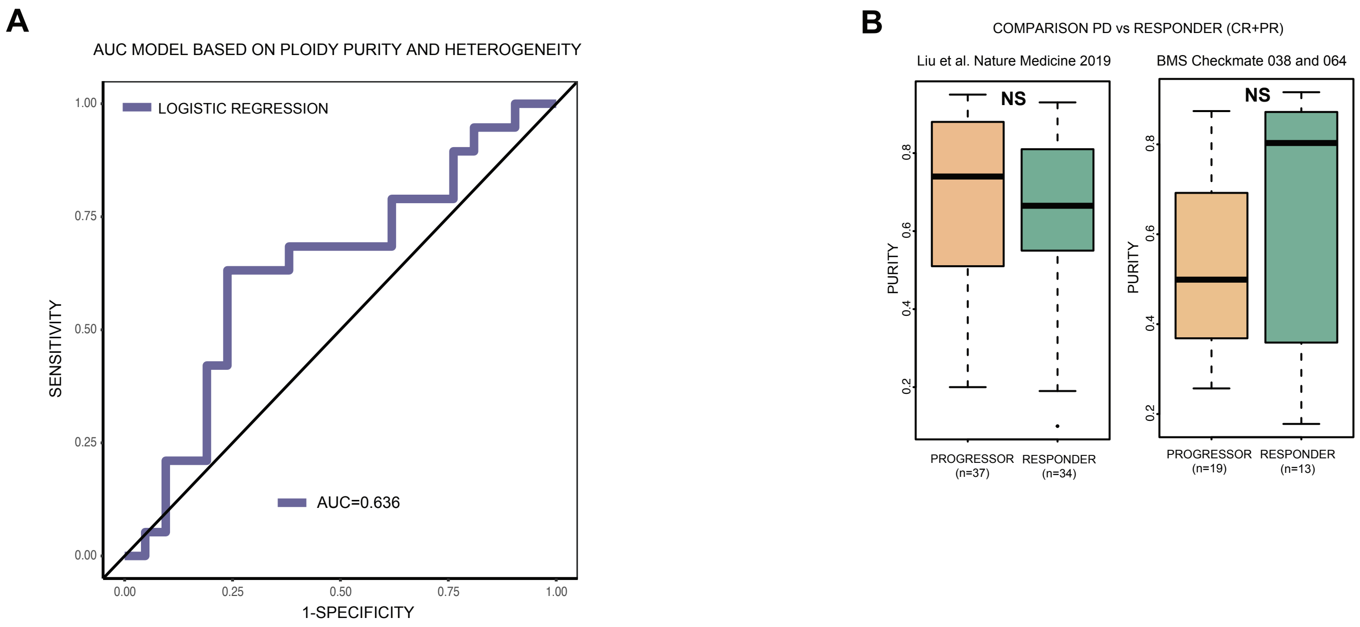

**Supplementary Figure1. Evaluation of previously nominated predictive model utilizing genomic heterogeneity, ploidy, and purity in validation cohort. A.** ROC curve of the model implemented for ipi-naïve patient from the Liu et al. Nature Medicine 2019 paper. **B.** Tumor purity in responders (CR/PR as best response, green) vs progressors (PD, orange) in the original discovery cohort (left) and the clinical trial validation cohort (right) (MWW $p = 0.19$, $p = 0.4$ in the original cohort and clinical trial validation cohort, respectively).

**Supplementary Figure 2. Performance of logistic regression and decision tree models predicting intrinsic PD-1 ICB resistance using genomic heterogeneity and ploidy in the new combined (original discovery + clinical trial validation) cohort**

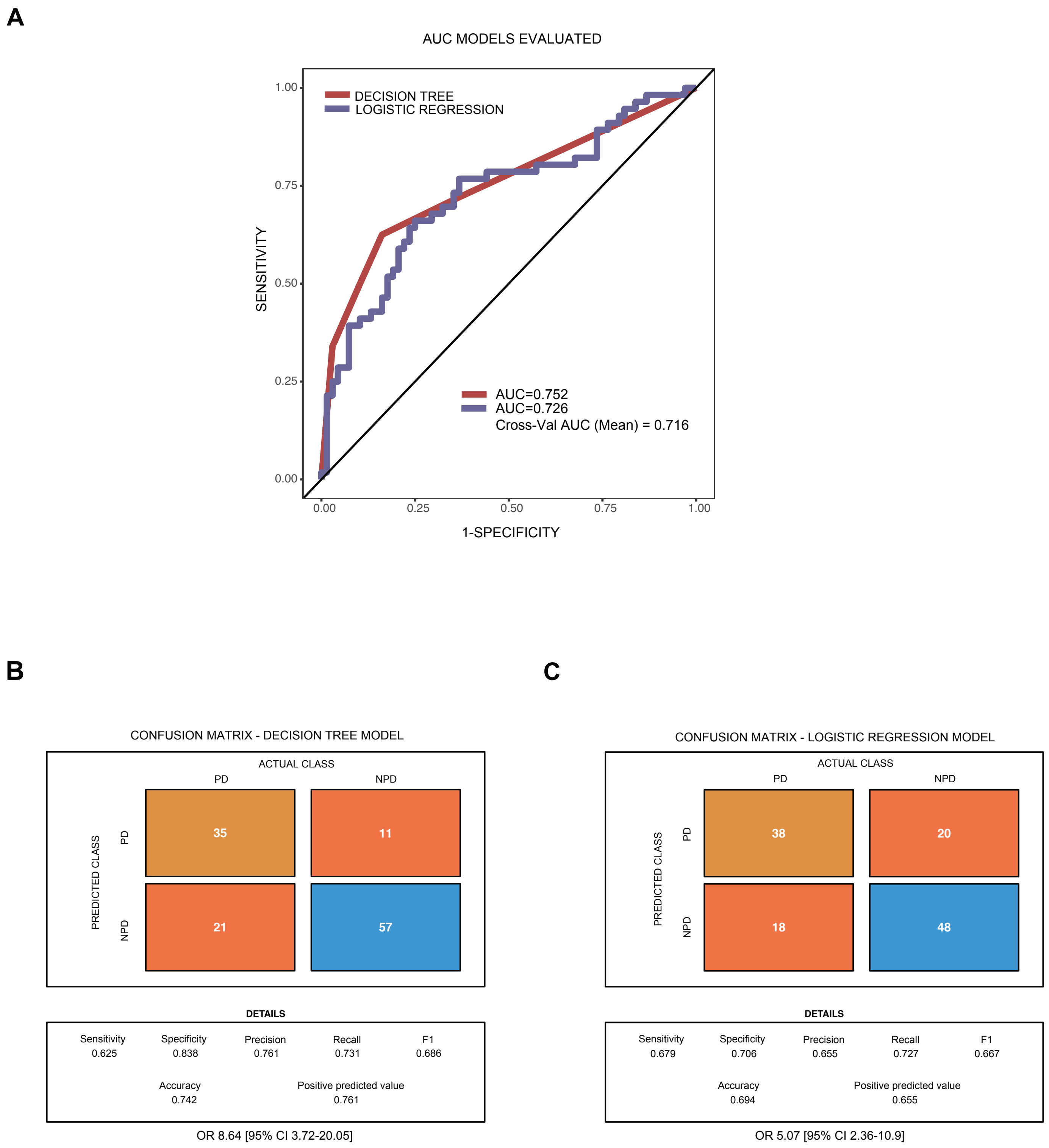

**Supplementary Figure2. Performance of logistic regression and decision tree models predicting intrinsic PD-1 ICB resistance using genomic heterogeneity and ploidy in the new combined (original discovery + clinical trial validation) cohort. A.** ROC curves of the two models implemented, decision tree (red) and logistic regression (purple). **B.** Confusion matrix for the decision tree model. **C.** Confusion matrix for the logistic regression model.

**Supplementary Figure 3. Survival stratification by model predictions.**

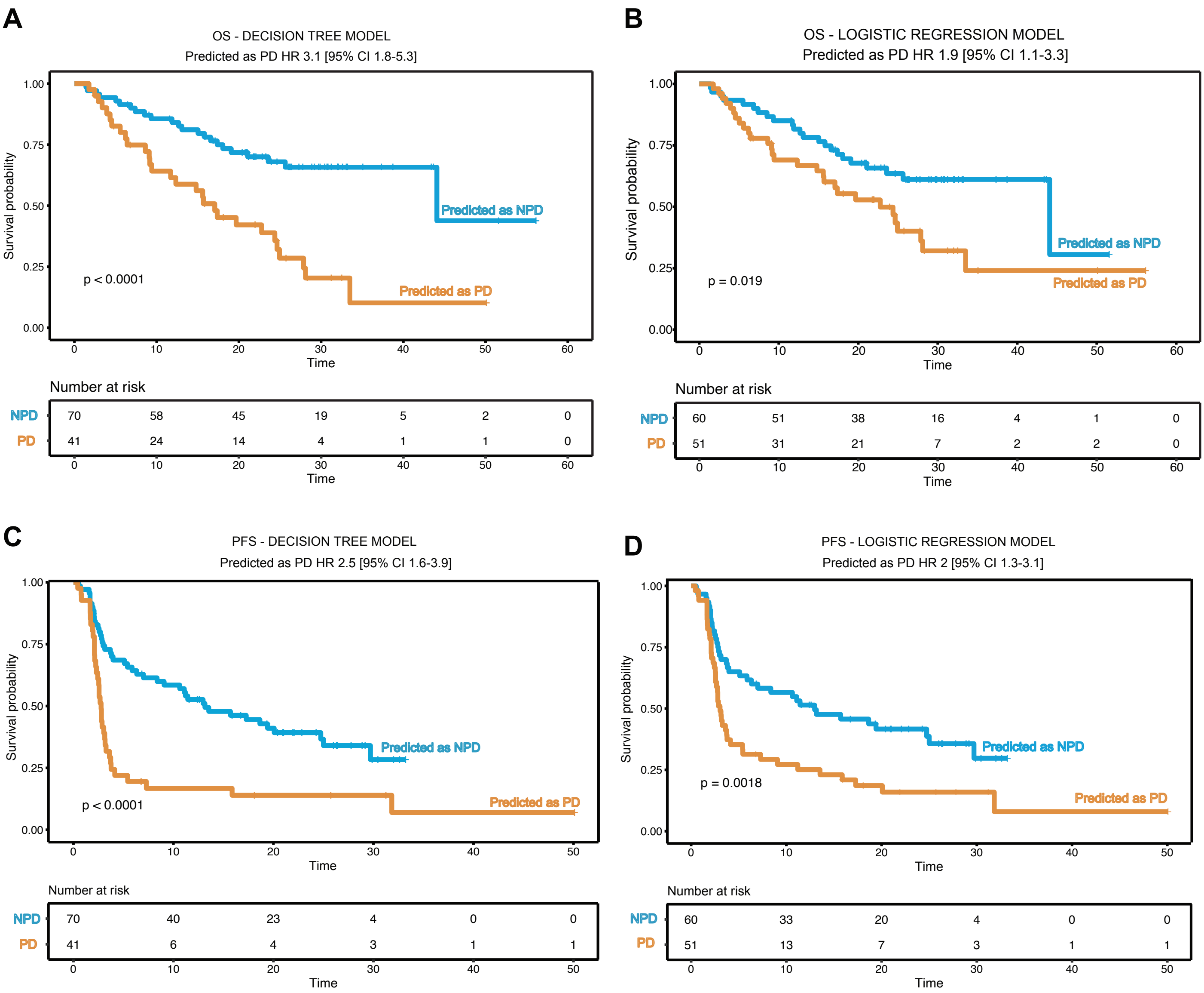

**Supplementary Figure3. Survival stratification by model predictions. A.** Overall survival curve stratified by patients predicted by the decision tree model as PD (orange) and NPD (blue) (log rank $p < 0.0001$). **B.** Overall survival curve stratified by patients predicted by the logistic regression model as PD (orange) and NPD (blue) (log rank $p = 0.019$). **C.** Progression free survival curve stratified by patients predicted by the decision tree model as PD (orange) and NPD (blue) (log-rank $p < 0.0001$). **D.** Progression free survival curve stratified by patients predicted by the logistic regression model as PD (orange) and NPD (blue) (log-rank $p =0.0018$).

**Supplementary Figure 4. Whole-genome doubling in responder vs progressor patients to PD-1 ICB.**

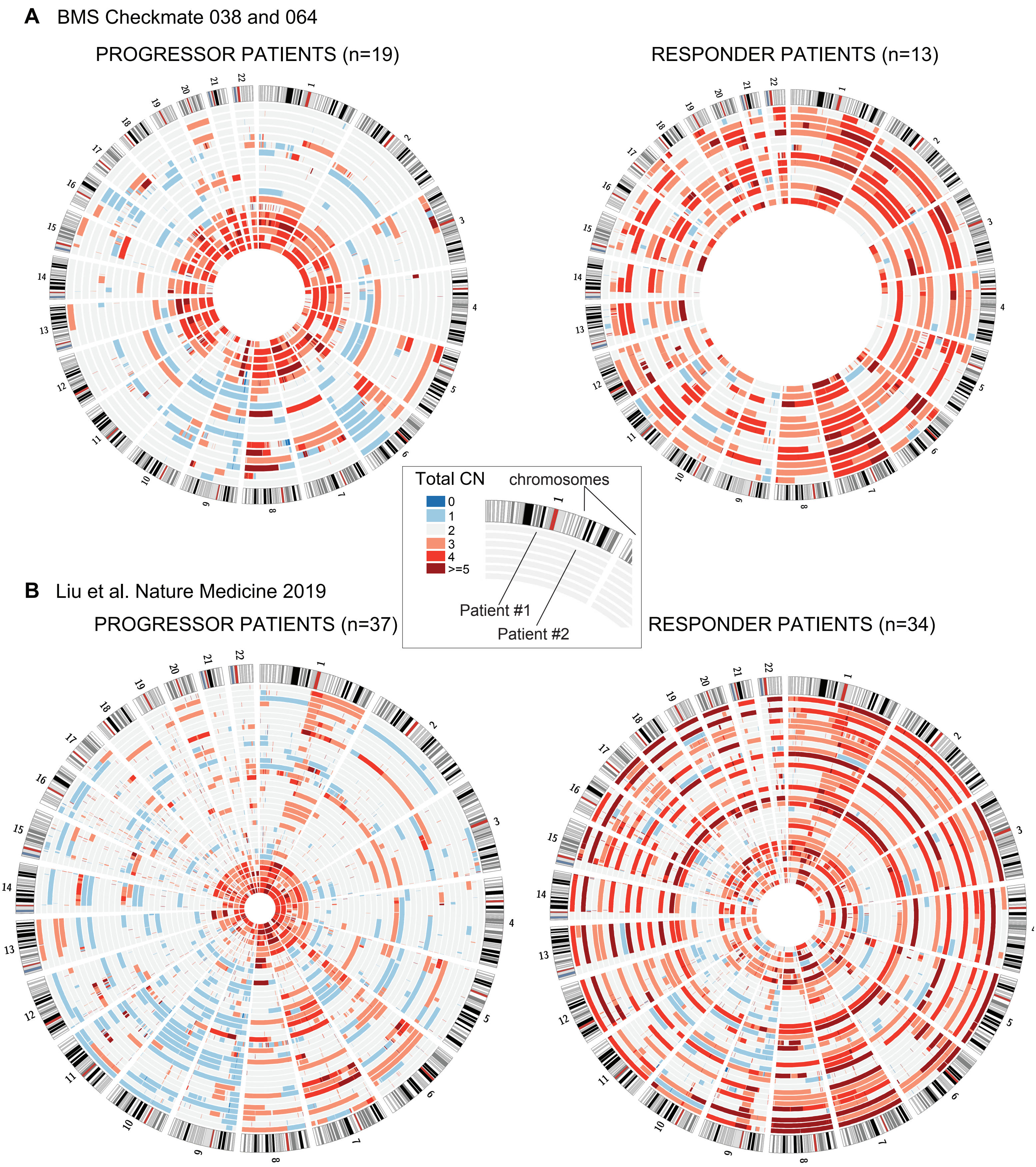

**Supplementary Figure4. Whole-genome doubling in responder vs progressor patients to PD-1 ICB.**

**A.** Circos plot of the total copy number across the genome (online methods) of progressors (PD as best response, left) and responders (CR/PR as best response, right) in the clinical trial validation cohort . Each circle represents a patient. Region of the chromosomes with a total copy number greater than 2 are showed in red; total copy number lower than 2 are showed in blue. **B.** Circos plot of the total copy number across the genome (online methods) of progressors (PD as best response, left) and responders (CR/PR as best response, right) in the original discovery cohort.

**Supplementary Figure 5. Supplementary Figure5. SNV multiplicity ratio and association with PD-1 ICB resistance.**

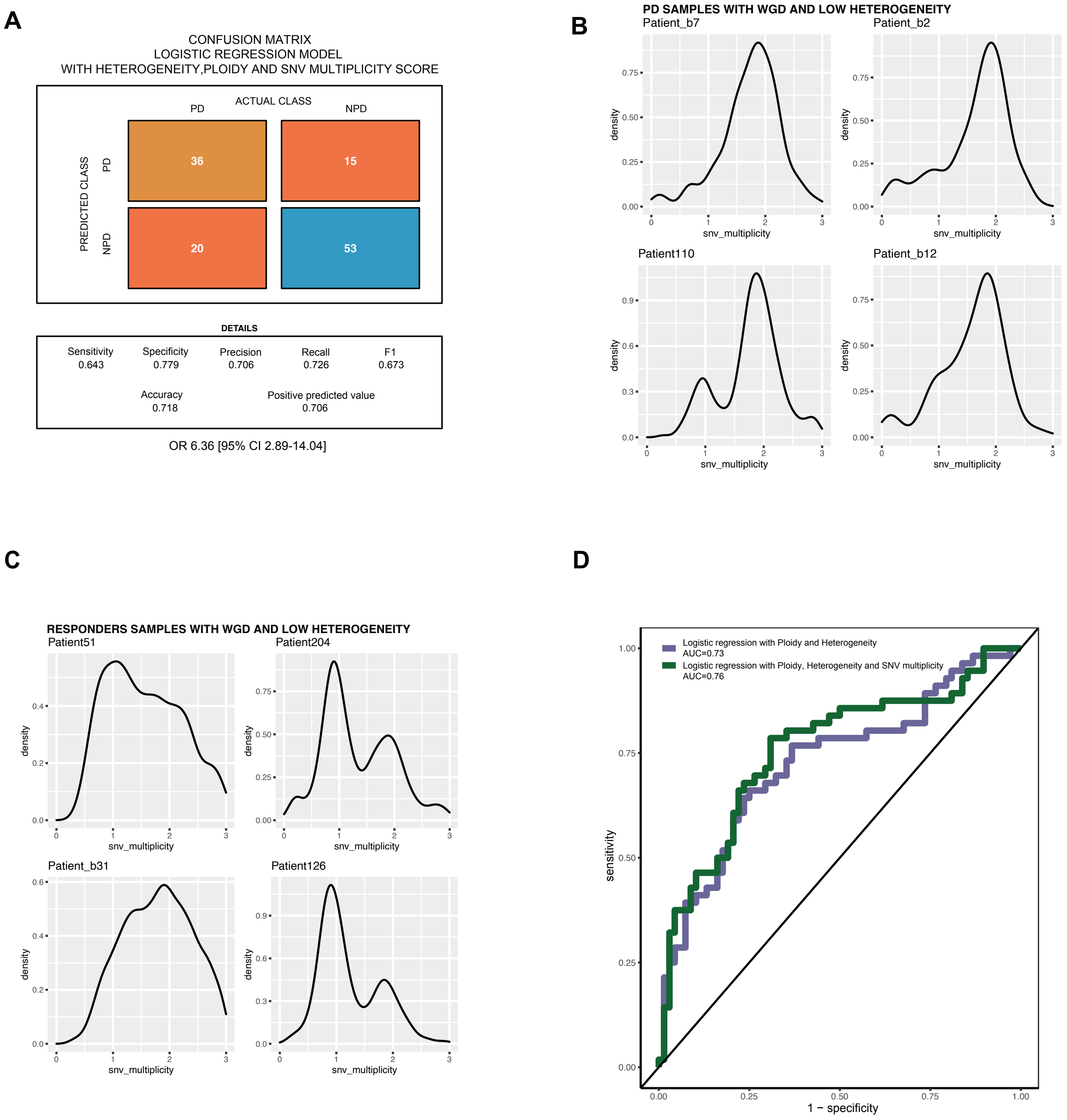

**Supplementary Figure5. SNV multiplicity ratio and association with PD-1 ICB resistance. A.** Confusion matrix of the logistic regression model trained using the three features (ploidy, heterogeneity and SNV multiplicity score). **B.** Distribution of multiplicity of mutations in misclassified (predicted NPD but observed PD) patient tumors with WGD and low heterogeneity **C.** Distribution of multiplicity of mutations in correctly classified (predicted and observed NPD) patient tumors with WGD and low heterogeneity. **D.** AUC of logistic regression models predicting PD with (green) and without (purple) including SNV multiplicity 2:1 ratio in addition to ploidy and heterogeneity.

**Supplementary Figure 6. Best treatment response by genomic heterogeneity, ploidy, and model prediction.**

A

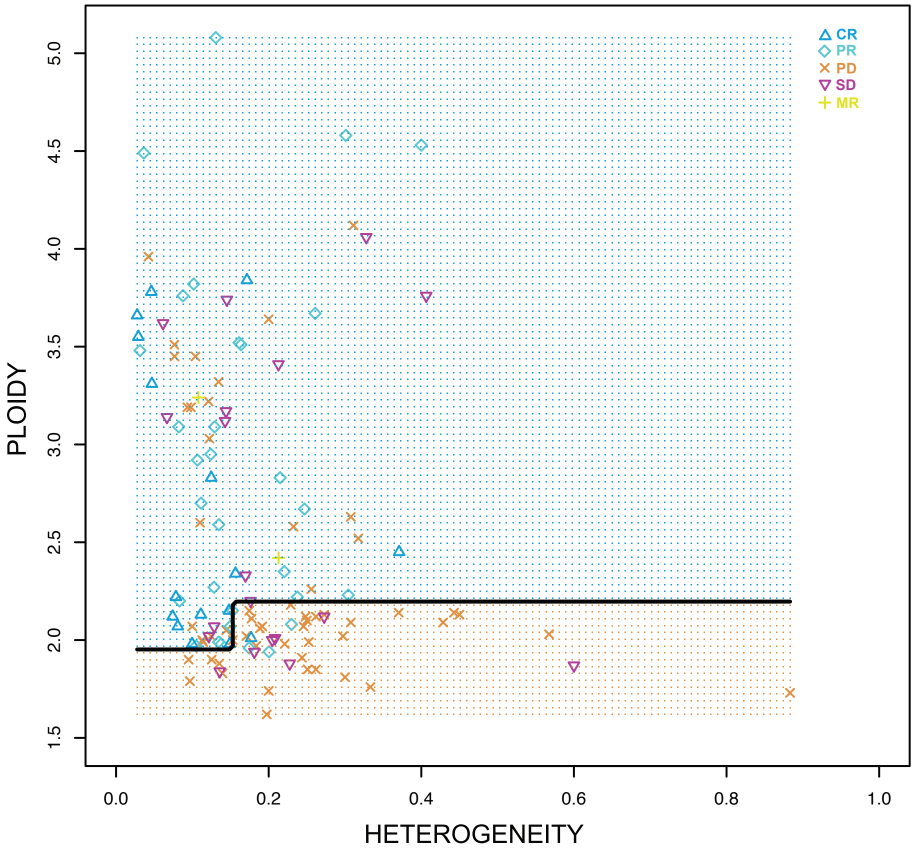


B

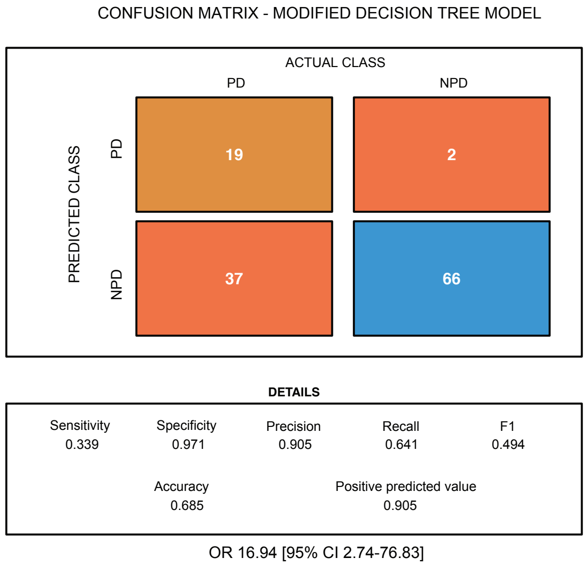


C

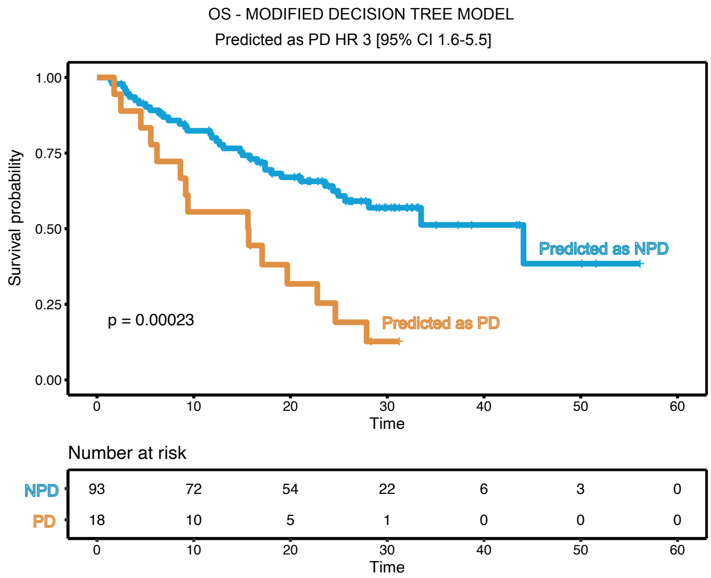

**Supplementary Figure6. Best treatment response by genomic heterogeneity, ploidy, and model prediction.** **A.** Decision boundaries of the Decision Tree model in which the symbol of each patient represents the best response. Most misclassifications in the predicted PD samples (orange area, bottom), occur at lower heterogeneity; above 0.25 the misclassified samples are NPD but best response is SD rather than CR/PR. **B.** Confusion matrix MDT; The model is characterized by high Precision and Specificity: 19/21 predicted PD patients are correctly predicted (90% precision/PPV), 66/68 patients with NPD are correctly predicted (97% specificity). **C.** Overall survival curve stratified by patients predicted by the MDT model as PD (orange) and NPD (blue) (log rank p = 0.00023); in the survival analysis have been excluded the samples from Checkmate 064 that received a sequential treatment (n=13).

**Supplementary Figure 7. Correlation of ploidy and heterogeneity with known clinical features.**

**A**

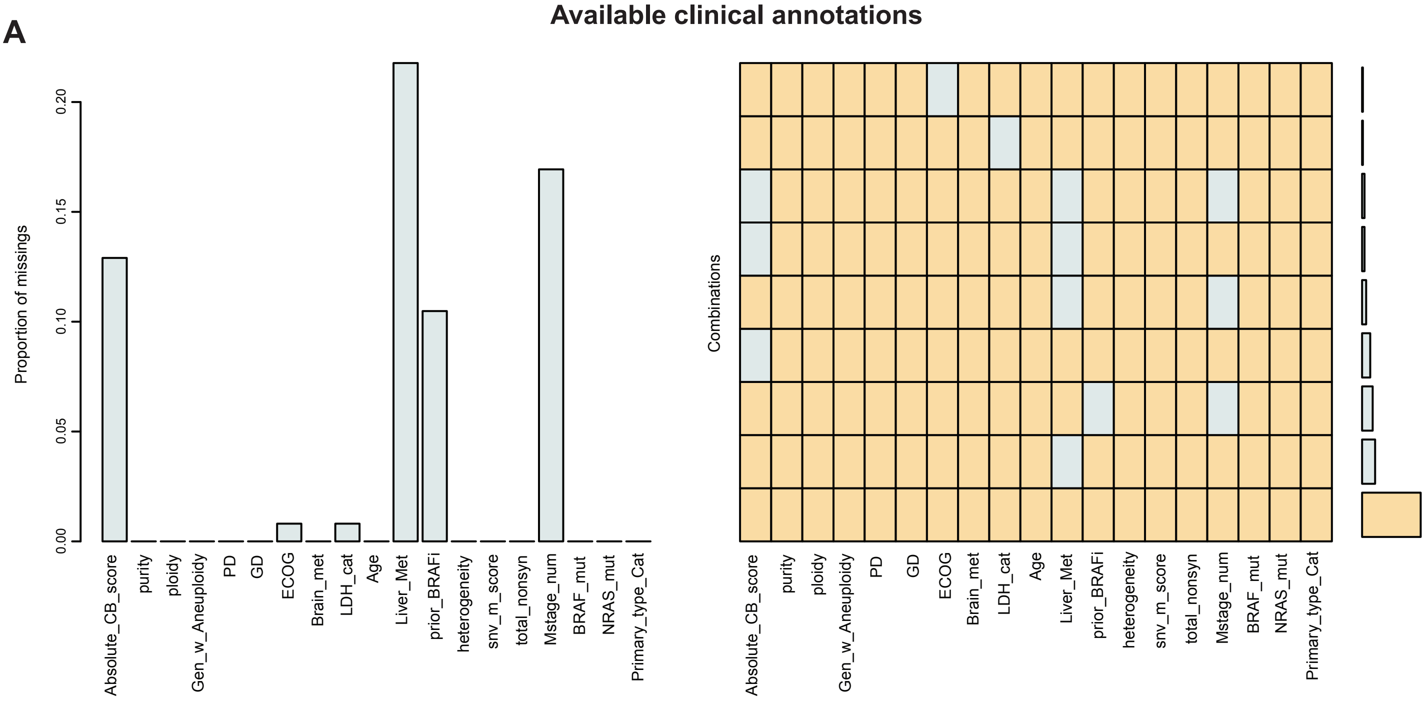


**B**

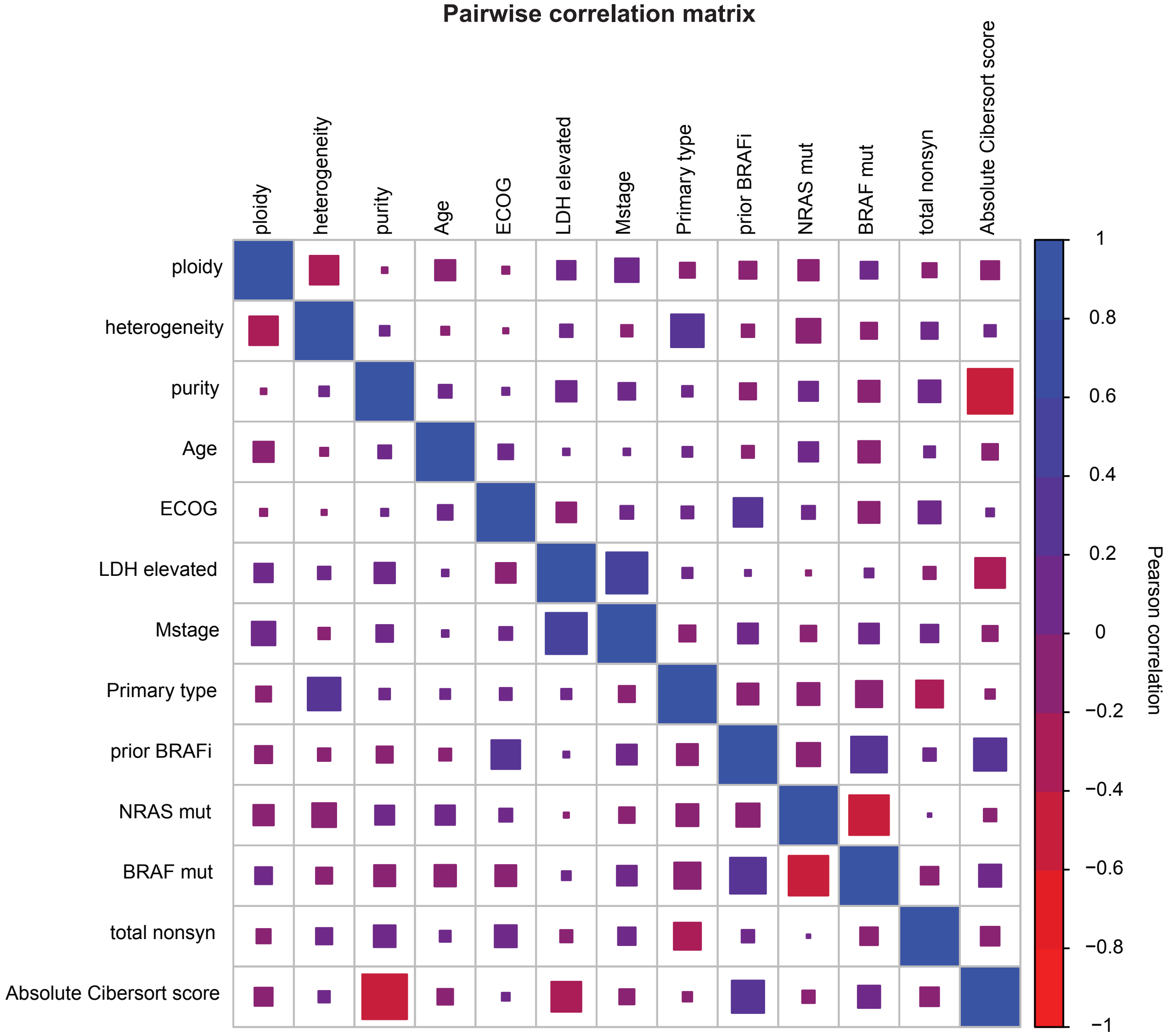

**Supplementary Figure7. Correlation of ploidy and heterogeneity with known clinical features.**
**A.** Proportion of missing values in the discovery cohort; The cibersortx quantification of the immune infiltrate was performed in the samples with available RNAseq; for some samples of the two BMS clinical trial the Mstage, the Liver Metastasis and the prior BRAF annotation is missing.
**B.** Pairwise correlation matrix between clinical and molecular variables.

**Supplementary Figure 8. Heterogeneity and ploidy in different primary melanoma subtypes.**

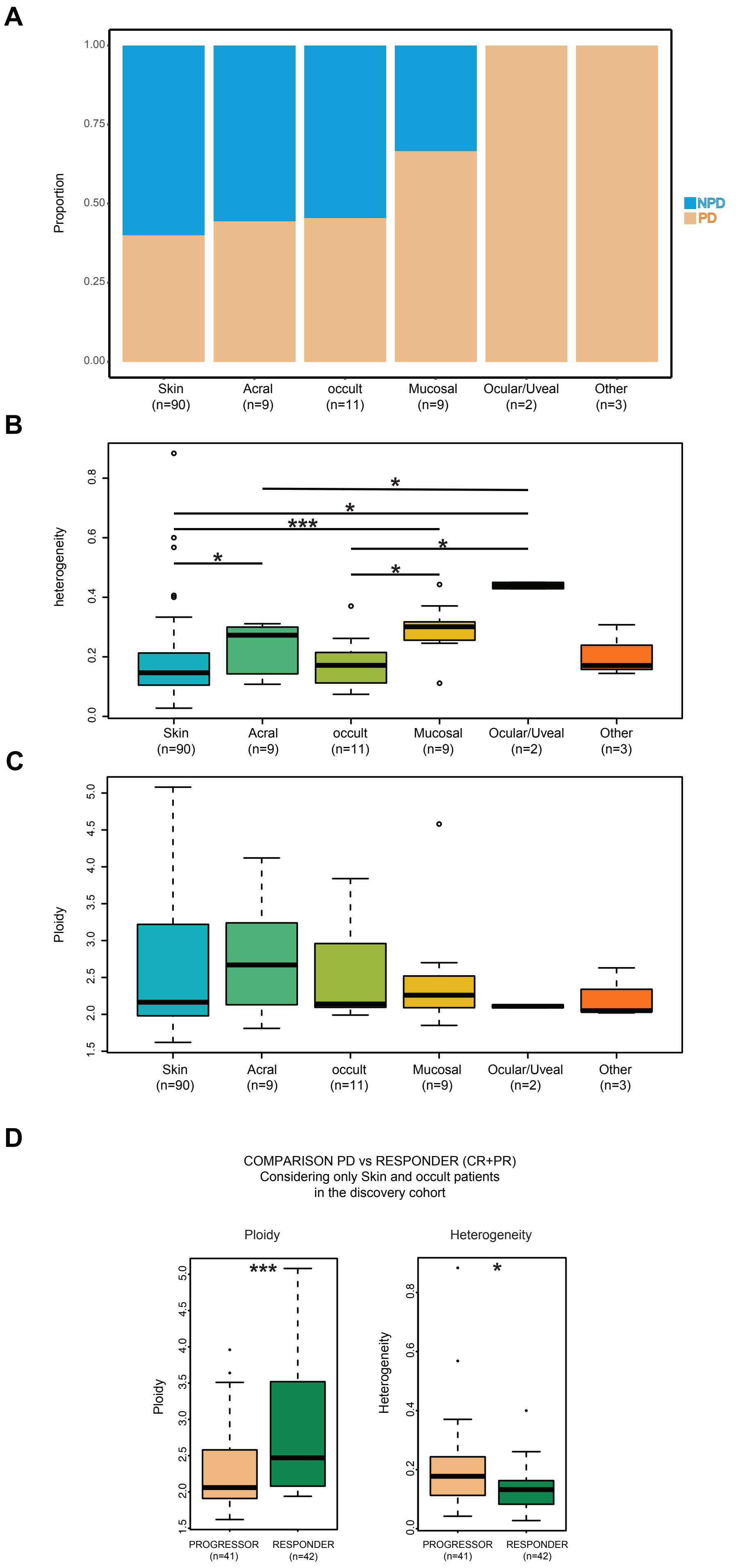

**Supplementary Figure8. Heterogeneity and ploidy in different primary melanoma subtypes. A.** Proportion of PD and NPD per primary type. **B.** Heterogeneity level across different Primary type. **C.** Ploidy level across different Primary type. **D.** Ploidy and heterogeneity in progressors (PD as best response) vs responders (CR/PR as best response) in the subcohort of cutaneous and occult melanomas.

**Supplementary Figure 9. Association of Mstage with heterogeneity and ploidy.**

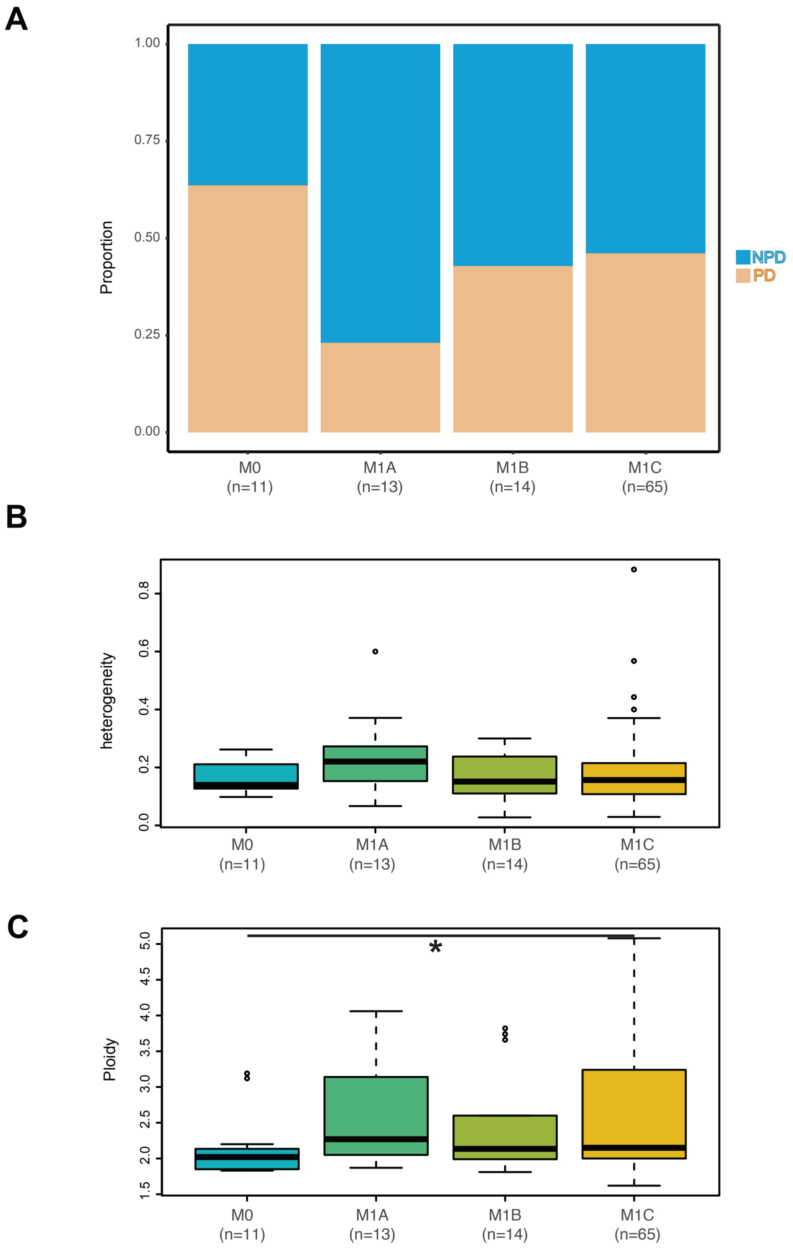

**Supplementary Figure9. Association of Mstage with heterogeneity and ploidy A.** Proportion of PD and NPD per mstage. **B.** Heterogeneity level across different mstage category. **C.** Ploidy level across different mstage category; significant stratification between M0 and M1C patients, with the M1C characterized by higher ploidy (MWW p = 0.032).

**Supplementary Figure 10. Association of brain metastasis and LDH levels with heterogeneity and ploidy**

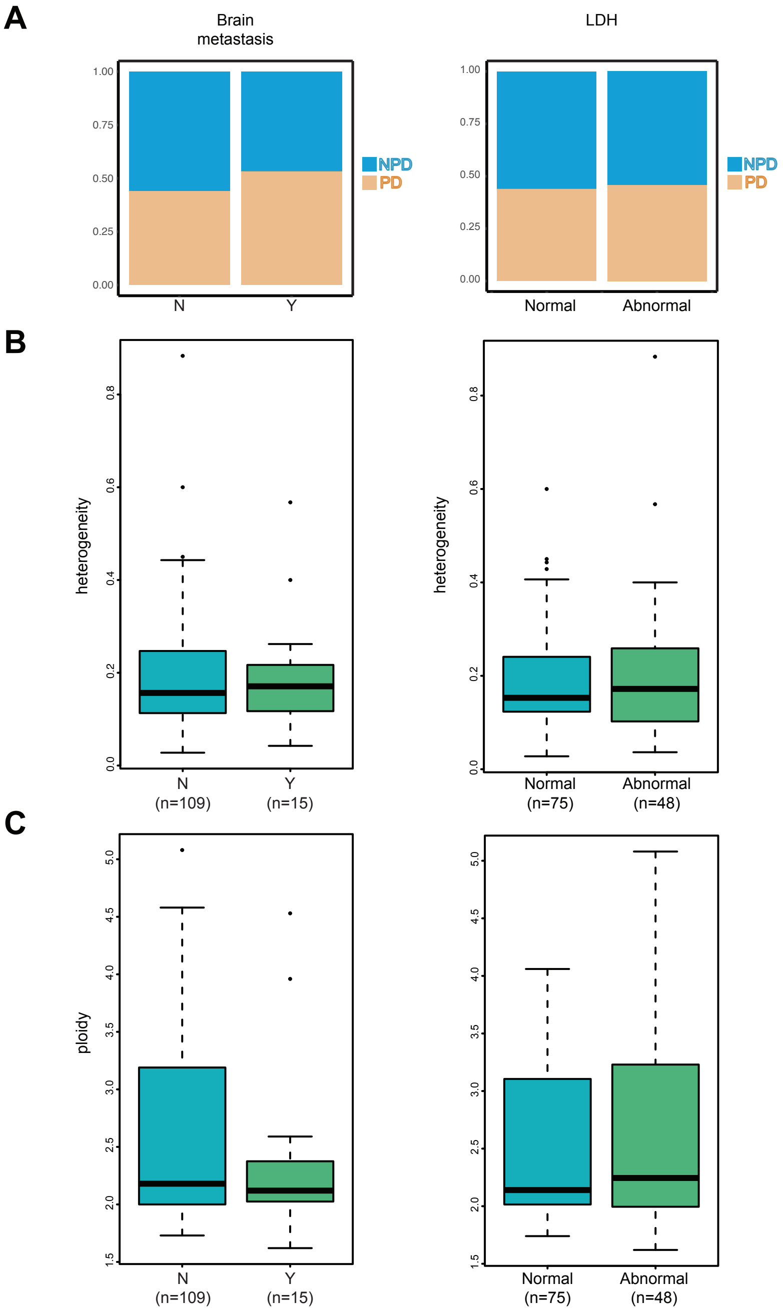

**Supplementary Figure10. Association of brain metastasis and LDH levels with heterogeneity and ploidy . A.** Proportion of PD and NPD per brain metastasis and LDH. **B.** Heterogeneity level comparison: patients with brain and without brain metastasis; patients with normal vs abnormal LDH level. **C.** Ploidy level comparison: patients with brain and without brain metastasis; patients with normal vs abnormal LDH level.

**Supplementary Figure 11. Evaluating TMB and response to PD-1 ICB with heterogeneity and ploidy.**

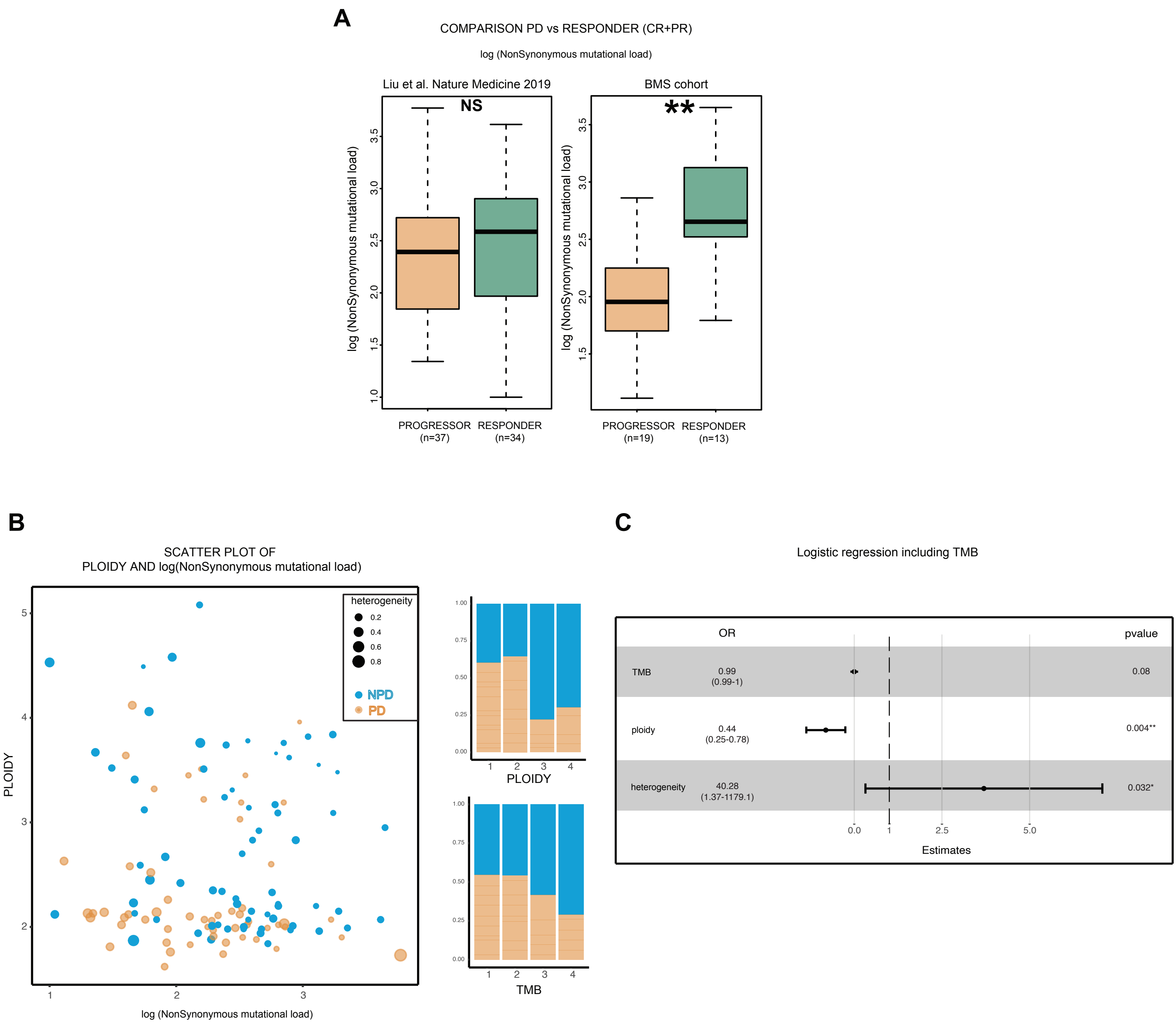

**Supplementary Figure11. Evaluating TMB and response to PD-1 ICB with heterogeneity and ploidy. A.** Tumor mutational burden (TMB) comparison between the extreme phenotypes patients in the discovery cohort, pvalue from wilcoxon rank-sum test. **B.** Scatterplot of ploidy and TMB, PD patients in orange and NPD patients in blue. The size of the dot represents the heterogeneity; the barplot on the right represent the proportion of PD and NPD patients in quartile groups according ploidy and TMB. **C.** Multivariate PFS COX regression model evaluation ploidy, heterogeneity and snv multiplicity score together with the TMB. Heterogeneity is a risk factor instead ploidy is a protective factor and they are independent from the TMB.

**Supplementary Figure 12.Cohort of patients with joint WES and RNAseq data from the combined discovery cohort.**

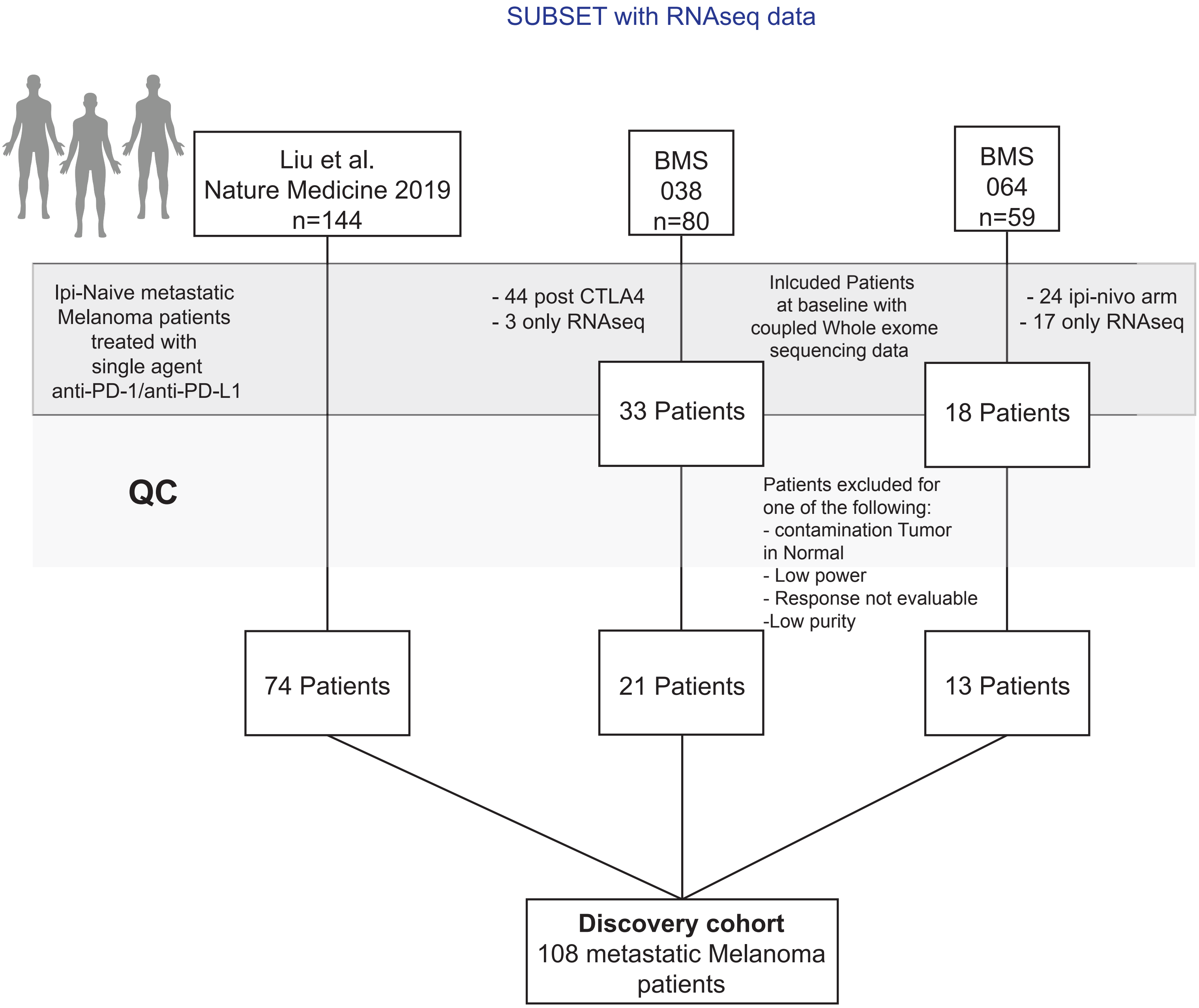

**Supplementary Figure12. Cohort of patients with joint WES and RNAseq data from the combined discovery cohort.** RNAseq data was available for a subgroup of samples (n=108).

**Supplementary Figure 13. Evaluating IFN-γ with heterogeneity and ploidy.**

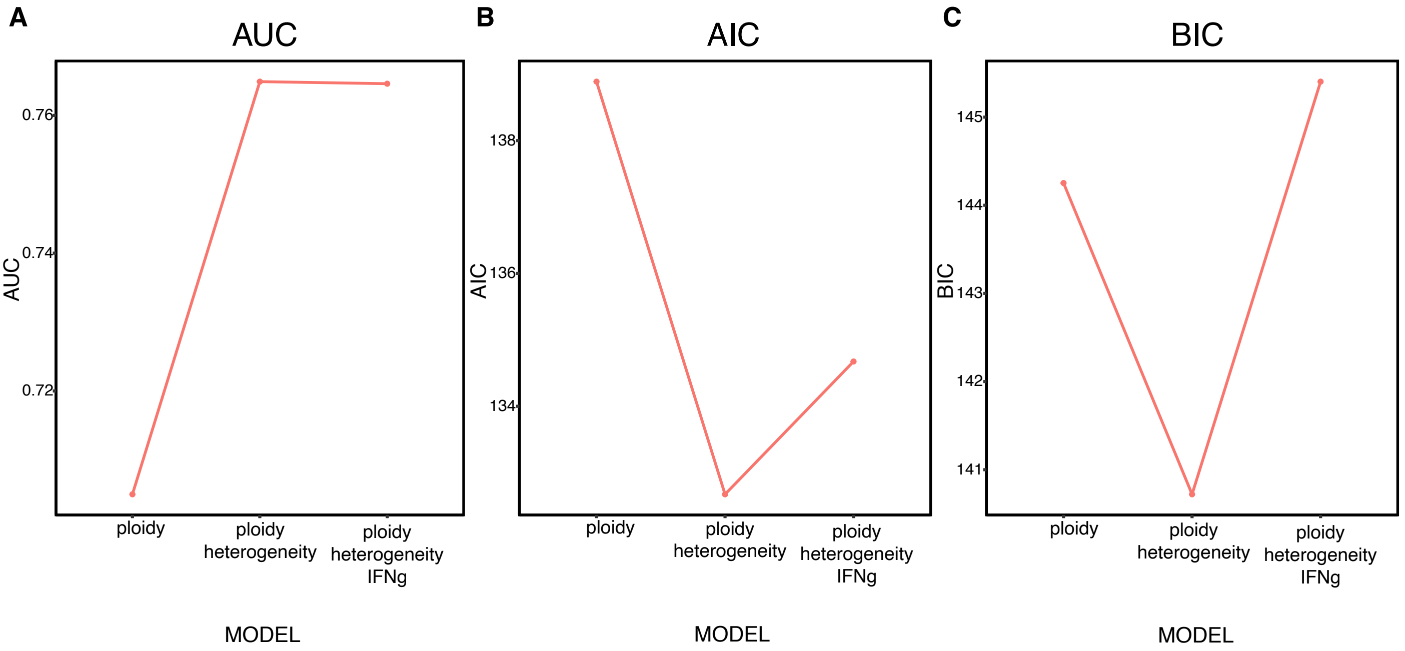

**Supplementary Figure13. Evaluating IFN-ɣ with heterogeneity and ploidy. A.** AUC for the logistic regression models evaluated. **B.** AIC for the models evaluated. **C.** BIC for the models evaluated.

**Supplementary Figure 14. Comparing a clinical nomogram to genomic model predictions.**

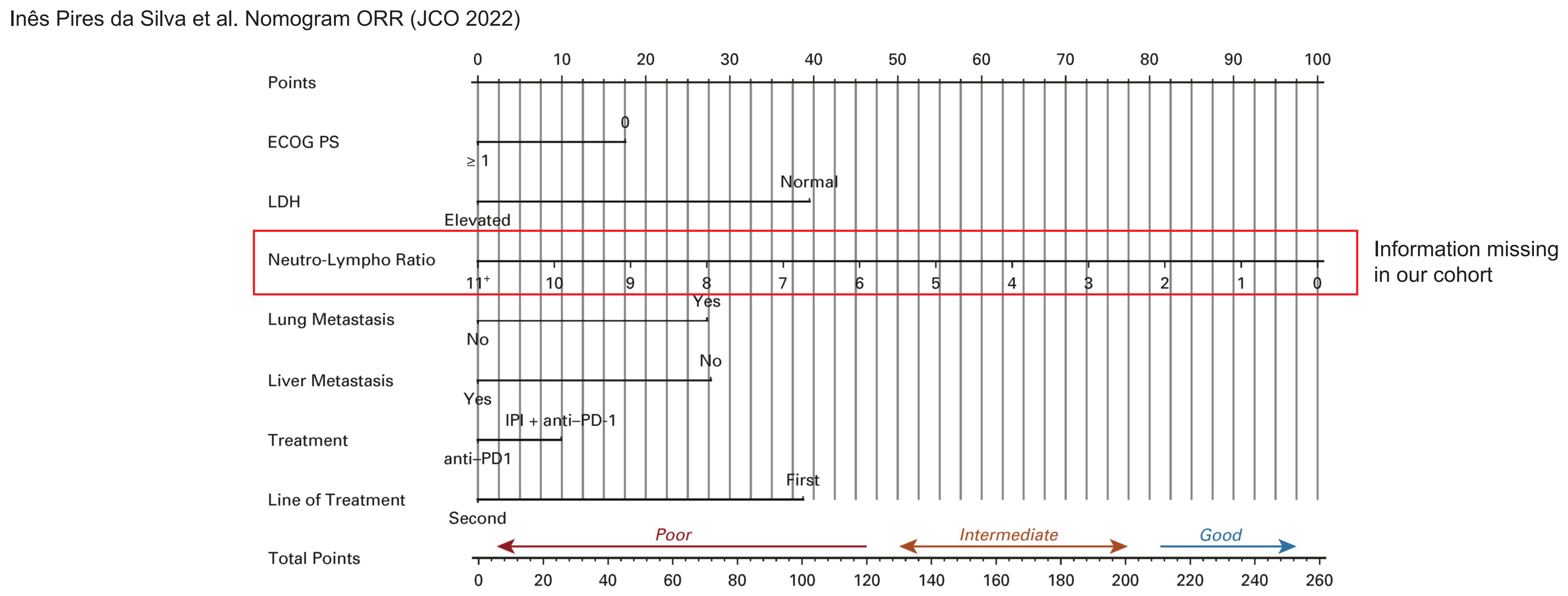


| | Sample_id | ECOG | LDH_cat | Liver_Met | Lung_Met | Line_of_treatment | Ploidy | Heterogeneity | BR | Nomogram score | Nomogram result | DT | modDT | LR |
|---|---|---|---|---|---|---|---|---|---|---|---|---|---|---|
| 4 | Patient14 | 0 | 0 | 0 | 1 | 0 | 1.81 | 0.3000000 | PD | 150 | intermediate or good | PD | PD | PD |
| 5 | Patient98 | 0 | 0 | 0 | 1 | 0 | 1.99 | 0.2525597 | PD | 150 | intermediate or good | PD | PD | PD |
| 6 | Patient134 | 0 | 0 | 0 | 1 | 0 | 2.12 | 0.2484277 | PD | 150 | intermediate or good | PD | PD | PD |
| 7 | Patient77 | 0 | 0 | 0 | 1 | 0 | 1.98 | 0.2209302 | PD | 150 | intermediate or good | PD | NPD | PD |
| 8 | Patient188 | 1 | 0 | 0 | 1 | 0 | 2.13 | 0.2727273 | PD | 132.5 | intermediate or good | PD | PD | PD |

**Supplementary Figure14. Comparing a clinical nomogram to genomic model predictions. A.** Patients predicted as poor or intermediate responder using the nomogram presented by Inês Pires da Silva et al. (DOI: 10.1200/JCO.21.01701 Journal of Clinical Oncology 40, no. 10 1068-1080.) that are good responders; the three models developed, except for one model for one sample, classify these samples as NPD due to the high ploidy and low heterogeneity. Since we don't have the NLR the score estimation is underestimate, for these samples however even with the maximum score of NLR will be classified as poor or intermediate from the nomogram.

**Supplementary Figure 15.Heterogeneity and Ploidy comparison between biopsy sites.**

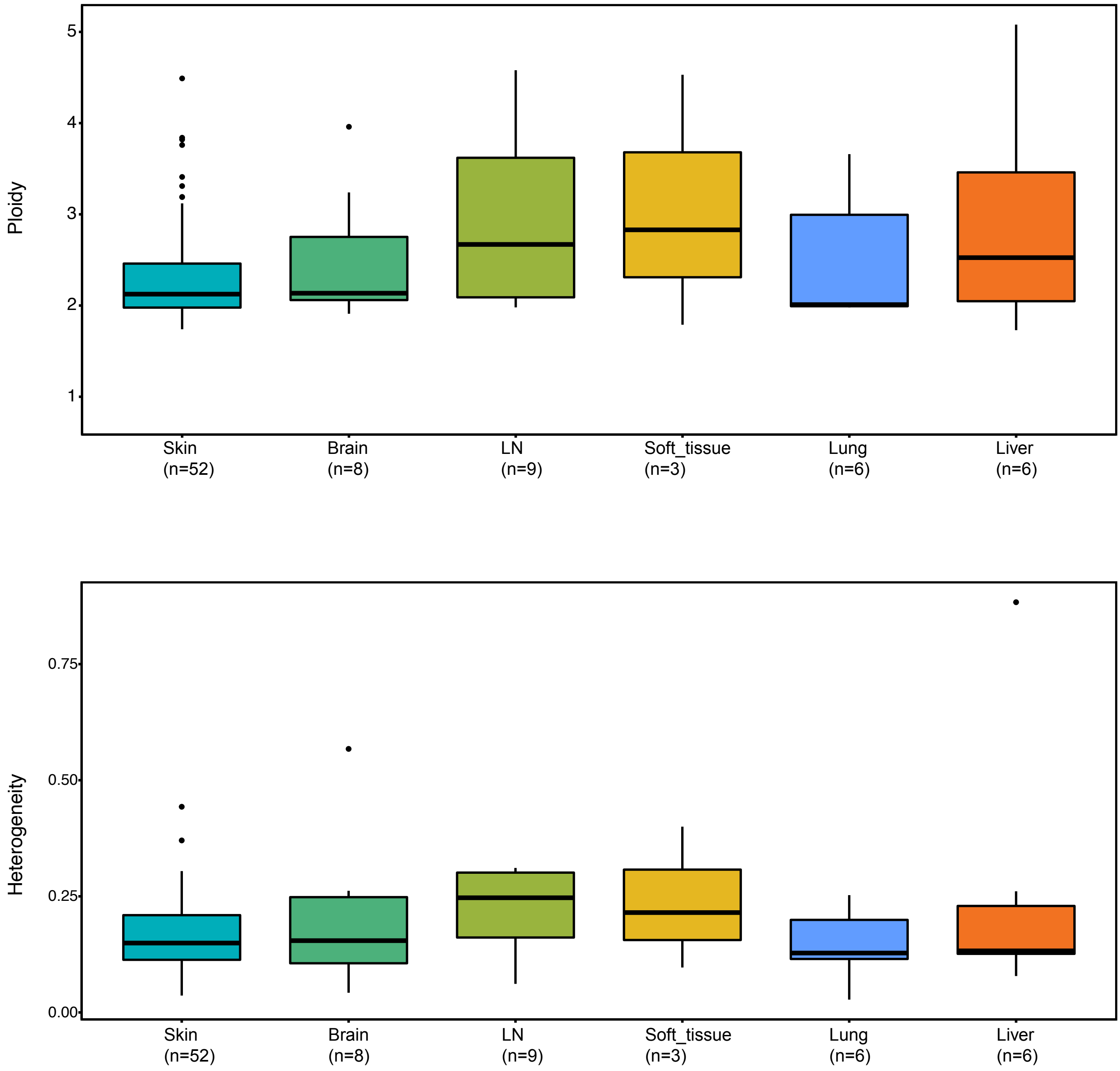

**Supplementary Figure15. Heterogeneity and Ploidy comparison between biopsy sites.**
Ploidy and Heterogeneity comparison between biopsy sites (annotation available only for the Liu et al. Nature Medicine 2019 cohort (n=84).

**Supplementary Figure 16. Heterogeneity and Ploidy association with ICB response using an automated heterogeneity and ploidy caller.**

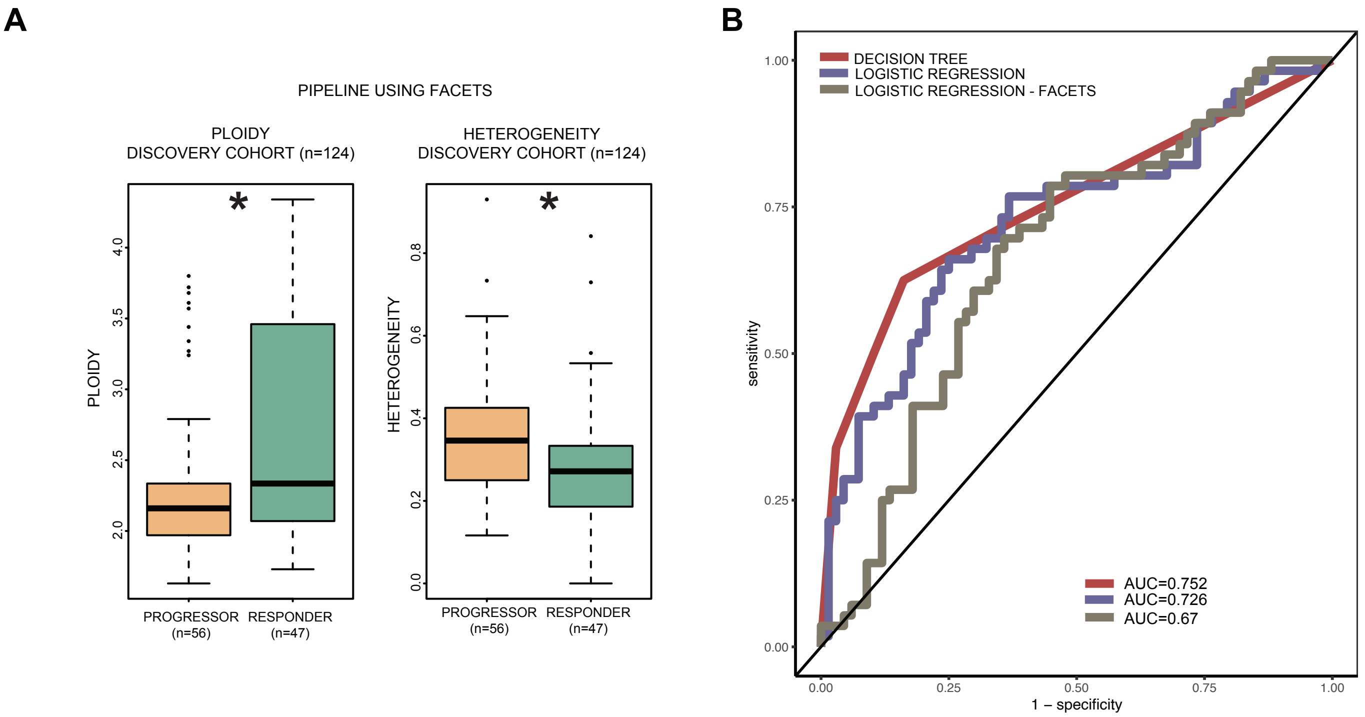

**Supplementary Figure16. Heterogeneity and Ploidy association with ICB response using an automated heterogeneity and ploidy caller.** Ploidy and Heterogeneity comparison between Responders (CR+PR) and Progressor using FACETS for Ploidy, Purity and cancer cell fraction estimation.

**Supplementary Figure 17. PPV, sensitivity and specificity values from the 4 fold CV procedure to identify the relative weight to use in the modified decision tree**

A

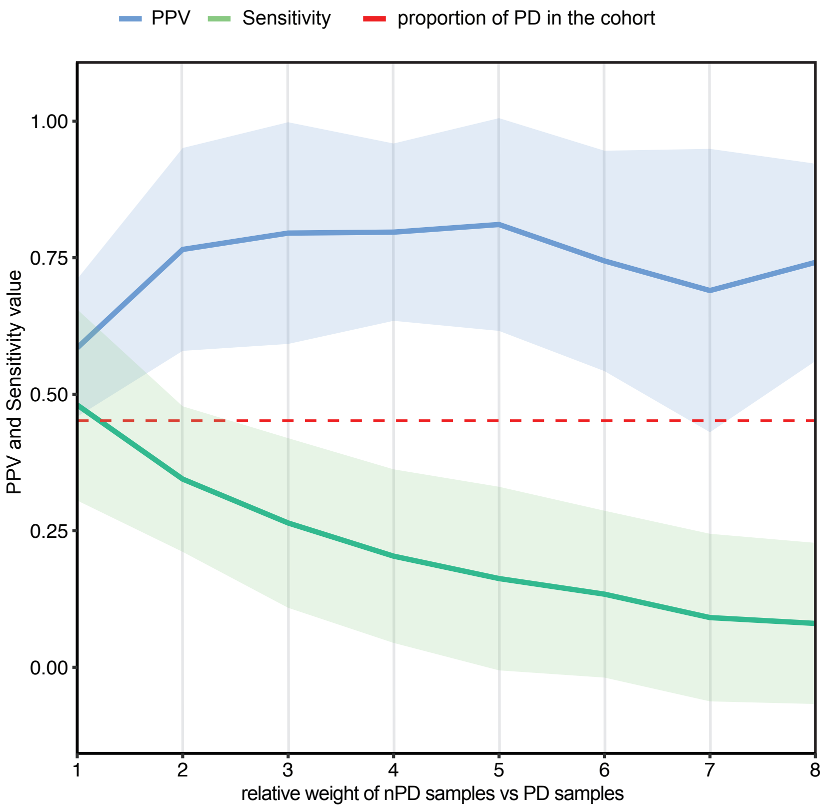


B

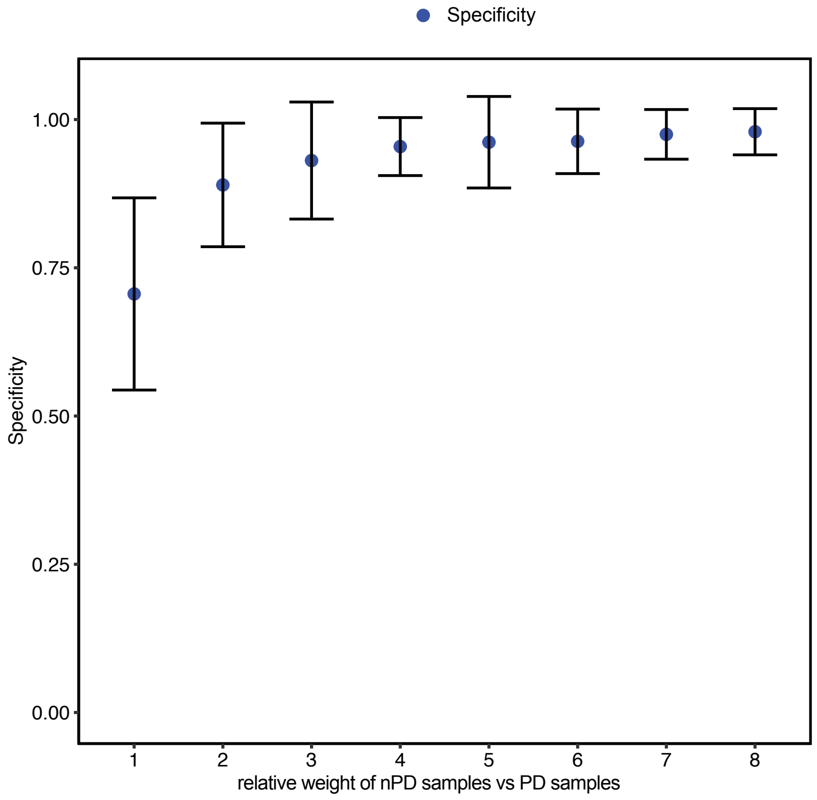

**Supplementary Figure17. PPV, sensitivity and specificity values from the 4 fold CV procedure to identify the relative weight to use in the modified decision tree.** The modified decision tree model to optimize precision was obtained by increasing relative weight of nPD samples vs PD samples in a 4-fold cross validation procedure repeated 10 times (using R package caret v 6.0.93). The choice of relative weight (nPD = 2) for the final model was selected for a tradeoff between increased precision or PPV (elbow method) and decreased sensitivity
